# A first-in-*Plasmodium* study on tRNA intron splicing endonuclease ‘*Pf*TSEN1’ and its substrate expression in clinical stage malaria

**DOI:** 10.1101/2024.06.07.597859

**Authors:** Mukesh Kumar Maurya, Ankita Behl, Amandeep Kaur Kahlon, Fernando De Leon, Palak Middha, Reena Nirban, Prerna Joshi, Jhalak Singhal, Geeta Kumari, Akshay Munjal, Rumaisha Shoaib, Neha Jha, Jagriti Pandey, Tanmay Dutta, Christoph Arenz, Anand Ranganathan, Shailja Singh

## Abstract

Mature tRNAs play critical role in several cellular processes including protein translation, post-translational-modifications and programmed-cell-death. Maturation of pre-tRNAs require removal of 5’-leaders, 3’-trailers, splicing of introns and addition of conserved 3’-terminal CCA sequence. The tRNA splicing mechanism, an essential step in tRNA maturation govern by a tRNA splicing endonuclease. While the existence of functional tRNA splicing endonuclease(s) in *Plasmodium falciparum* has not been identified, its significance in other eukaryotes suggests a potential role in tRNA splicing event. Our study identified total tRNAs in *Plasmodium* and characterize a *Pf*tRNA splicing endonuclease (annotated as *Pf*TSEN1) recognised recently as a component of ribonucleoprotein (RNP) complex, and synthesized a naphthoquinone derivative as a novel anti-malarial compound (‘TSEN_i_’) targeting the functional activity of this protein. Enzyme activity assays elucidated that *Pf*TSEN1 catalyses splicing of *in vitro* transcribed pre-tRNA^leu^, the expression of which was confirmed during the clinical stages of malaria parasite by RT-PCR. Interestingly, TSEN_i_ binds to and inhibits enzymatic activity of *Pf*TSEN1, and showed potent anti-malarial activity against chloroquine-sensitive 3D7 and resistant strains Dd2 of *P. falciparum*. Overall, our study deliver key knowledge towards the functional role of *Pf*tRNA splicing endonuclease, and its inhibitor TSEN_i_ as potent anti-malarial.

## Introduction

World Health Organization (WHO) recommends artemisinin-based combination therapies as first-line treatments for falciparum malaria (1). However emergence and spread of artemisinin resistance has threatened the efforts made to control and eradicate this deadly disease (2). Artemisinin resistance is prevalent in Southeast Asia and defined clinically by delays in parasites clearance after treatment with artemisinin-based regimens. Spread of artemisinin resistance to other malaria endemic regions particularly sub-Saharan Africa is of great concern owing to devastating outcomes, and makes the disease management extremely challenging (2, 3). The rapidity with which drug resistant is occurring demands the identification and characterization of novel drug targets that can lead to the development of anti-malarials. A study by Hollin et. al. performed proteome-wide studies and provided the core set of components of Ribonucleoprotein (RNP) complexes in *P. falciparum* (4). These complexes are assemblies of RNA molecules and proteins, and are involved in regulating post-transcriptional events of gene expression including splicing, stability, translation and degradation (5–7). One class of proteins that were found to be associated with RNP complexes were tRNA splicing endonucleases that play crucial role in mechanism of transfer RNA (tRNA) maturation, and that ultimately leads to protein translation.

Functional tRNA splicing endonucleases are essential enzymes found in all domains of life, and are involved in the maturation of intron-containing pre-tRNAs. The catalytic process involved in splicing is conserved between archaea and eukaryotes where a TSEN separates the exons from the intron and subsequently a tRNA ligase unites the exon region (8). The human TSEN and archaeal TSENs share a conserved catalytic triad of Tyr, His, and Lys residues which is responsible for cleaving the phosphodiester backbone (9). Although human and archaeal TSENs depict similarities, they follow different mechanisms of recognizing substrates. Archaeal enzymes identify a pseudosymmetric local secondary RNA structure called the bulge-helix-bulge (BHB) motif which is comprised of four nucleotide helix flanked by two bulges, each composed of three nucleotides (9, 10). This BHB motif is generally present in the tRNA anticodon arm but can also be present at other sites in the tRNA molecule. On the contrary, human TSENs identify asymmetric tertiary structure of the tRNA and here precursor tRNAs harbor a single intron placed exactly between the first and second nucleotide downstream of the anticodon called as canonical position (37/38) (9, 11,12). This critical base pair between the anticodon loop and intron is called as A-I base pair. Additionally, all archaeal enzymes form complexes as homodimers (α_2,_ ε_2_) or homotetramers (α_4_) or heterotetramers (α_2_β_2_) or whereas human TSEN enzyme exist as an asymmetric heterotetramer (αβγδ) (9). The simplest archaeal composition of a TSEN (e.g., Methanocaldococcus jannaschii) revealed that is a homotetramer (α4) with two subunits adopting a catalytically active conformation and two executing non-catalytic, structural roles. On the contrary eukaryotic TSEN complexes are comprised of four different subunits termed as TSEN2, TSEN34, TSEN54 and TSEN15, where TSEN2 and TSEN34 forms the catalytic subunits responsible for cleaving the 5’ and 3’ splice sites respectively and TSEN15 and TSEN54 are non-catalytic structural proteins whose defined role in splicing is not well characterized (9).

Despite the conventional roles of tRNAs in protein translation, recent reports indicate their key roles in multiple other processes, including post-translational modifications, programmed cell death, modulating global gene expression in response to stress, and disease progression (13). Considering the important functions of tRNAs in regulating multiple processes in normal physiology and disease, we for the first time attempted to identify total tRNAs and tRNA splicing endonucleases in *Plasmodium* (*Pf*TSENs), the key enzymes responsible for tRNA maturation, and delineated the first evidence for their functional relevance in malaria biology. Our *in silico* data mining revealed the existence of 78 tRNAs and two splicing endonucleases (*Pf*TSEN1 and *Pf*TSEN2) in *Plasmodium*. Structure-function characterization of *Pf*TSEN1 depicted its similarity with archaeal TSENs. *In vitro* enzymatic assays illustrated the ability of *Pf*TSEN1 to splice intron containing pre-tRNA^Leu^ (apicoplast) that marks the hallmark feature of a splicing endonuclease enzyme. Considering the essentiality and functional importance of *Pf*TSENs in parasite biology, we identified a RNA binding protein inhibitor ‘1-(3-aminopropyl)-1H-pyrrolo[2,3-f]quinoline-4,9-dione’ (TSEN_i_) that depicted binding with *Pf*TSEN1 and blocks its splicing endonuclease activity. Interestingly, TSEN_i_ showed potent anti-malarial activity against asexual blood stages of *Pf*3D7 and chloroquine drug-resistant strain *Pf*Dd2. Overall, our study contribute towards understanding the role of a novel splicing endonuclease of *Plasmodium* and decipher a small molecule TSEN_i_ targeting this protein and elucidating anti-malarial properties. This piece of knowledge can help in bridging the key knowledge gaps required for anti-malarial drug development and tackling the problem of artemisinin resistance.

## Results

### Identification of tRNAs in *Plasmodium* reveal the presence of a single intron containing pre-tRNA specific for leucine

We searched for the presence of pre-tRNAs in *Plasmodium* responsible for bringing the amino acids to ribosomes for protein production. Our search in Plasmodb database (14) and tRNAscan-SE 2.0 webserver (15) suggested the existence of 45 tRNAs in nucleus and 33 tRNAs in apicoplast (Table S1, S2). Out of these, 5 tRNAs specific for leucine were found in nucleus and 3 leucine specific tRNAs were identified in apicoplast. Detailed sequence analysis revealed that only one leucine specific tRNA (PF3D7_API00300, 220 bases) contains 135 bases long intron (37 -171) that is encoded by apicoplast genome [annotated as pre-tRNA^Leu^ (apicoplast)] (Table S1). Additionally, tRNAscan-SE 2.0 webserver identified one tyrosine specific tRNA (PF3D7_0702800, 73 bases) of nuclear genome to harbour 11 bases long intron (38-48), however this intronic region was not identified in Plasmodb database. 2D structures of pre and mature tRNA^Leu^ (apicoplast) were modelled using RNAfold webserver (16) that clearly demonstrate the presence of a long intron (37-171 nucleotides), that is removed to restore the anti-codon (UAA) for leucine (Fig. 1A i, ii). 2D structure of tRNA^Leu^ (nucleus) without intron is shown as reference in fig. 1 A iii.

**Figure 1:**
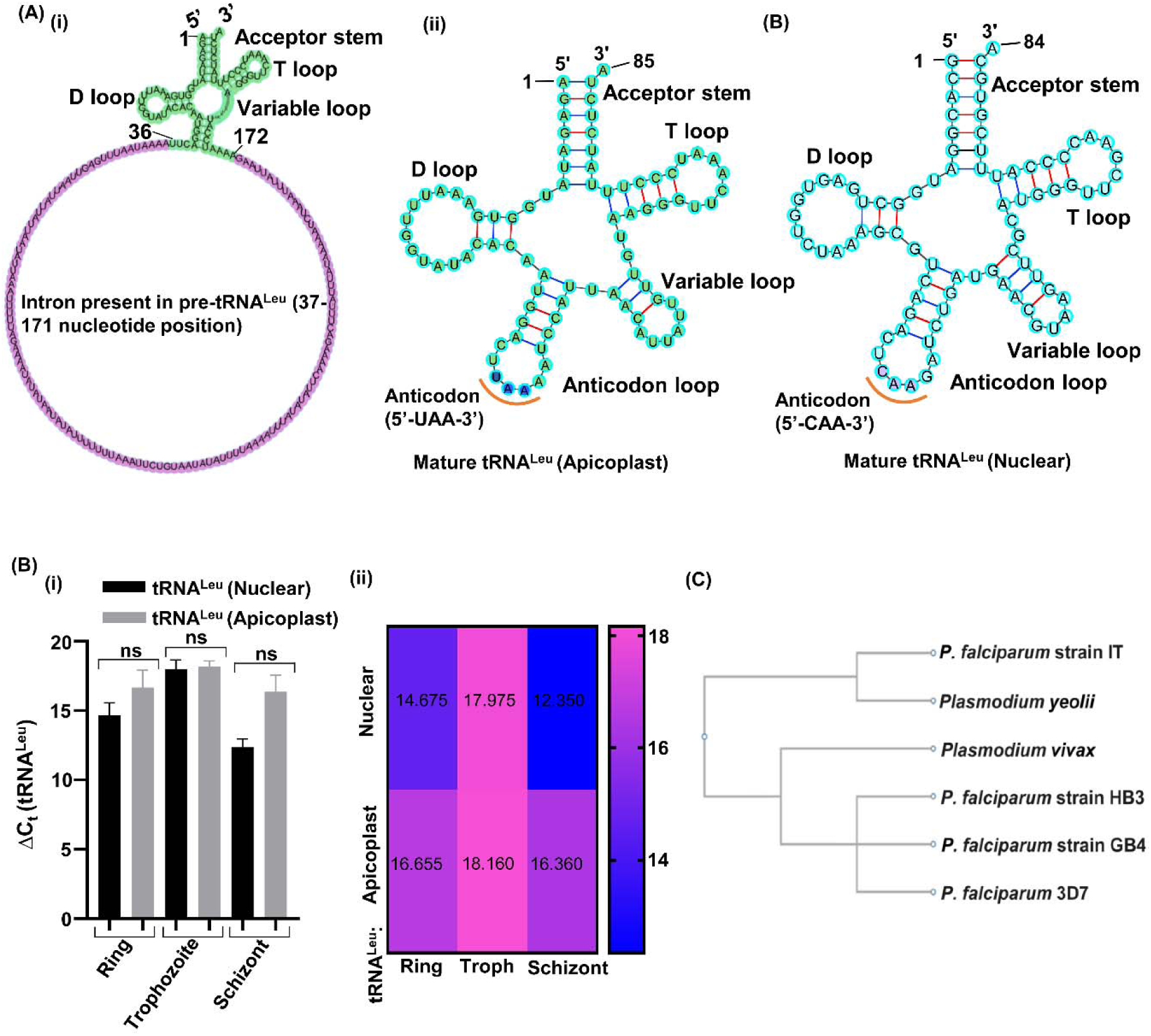
*In silico* analysis of *Plasmodium* tRNAs. **(A i)** 2D structure of pre-tRNA^Leu^ (apicoplast) in its cloverleaf model. **(ii)** 2D structure of mature tRNA^Leu^ (apicoplast) in its cloverleaf model. Removal of intron restores the anticodon (UAA) as depicted. (iii) 2D structure of tRNA^Leu^ (nucleus) with no intronic region is shown as reference. **(B i)** Relative expression of tRNA^Leu^ (Apicoplast) and tRNA^Leu^ (nuclear) at transcript levels during asexual blood stages of malaria parasite using qRT-PCR. Error bars represent standard deviation among three replicates. ns represents non-significant difference. **(ii)** Heat map depicting differential expression of tRNA^Leu^ (Apicoplast) and tRNA^Leu^ (nuclear) at different intraerythrocytic stages. **(C)** Phylogenetic tree demonstrating the evolutionary relationships of tRNA^Leu^ (Apicoplast) with its homologs in different *Plasmodium* strains (GB4, HB3, IT) and species (*Plasmodium vivax*, *Plasmodium Yeolii*.

We next analysed the expression of pre-tRNA^Leu^ (Apicoplast) and pre-tRNA^Leu^ (nucleus) during the asexual blood stages of *Pf*3D7. Transcript levels of pre-tRNA^Leu^ (Apicoplast) and pre-tRNA^Leu^ (nucleus) clearly demonstrated their expression throughout the intra-erythrocytic life cycle of *P. falciparum* (Fig. 1 B). Phylogenetic analysis of pre-tRNA^Leu^ (Apicoplast) with intron in different *Plasmodium* strains and species suggest the conserved nature of this tRNA (Fig. 1 C). Multiple sequence alignment of pre-tRNA^Leu^ (Apicoplast) with its homologs is shown in fig. S1.

### tRNA spicing endonucleases harboring a conserved catalytic domain identified in *Plasmodium falciparum* genome

We next looked for the presence of tRNA splicing endonucleases (TSENs) in *Plasmodium* that are required to cleave the intron from pre-tRNA^(Leu)^ (Apicoplast) to form mature tRNA. The TSEN complex is prerequisite for catalysis of tRNA splicing in Archaea and Eukarya. Compared to archaea and eukarya, no tRNA splicing endonucleases have been characterized in *Plasmodium* till date and their role in the life cycle of *Plasmodium falciparum* remains largely unknown. Therefore, the existence of tRNA spicing endonucleases in malaria parasite was identified and their functional relevance was explored through bioinformatics analysis. Uniprot (17) was used to search *P. falciparum* genome for presence of tRNA spicing endonuclease genes, and two genes were obtained (A0A144A1E9_PLAF7 and Q8I718_PLAF7), indicating the presence of tRNA spicing endonuclease in *Plasmodium* species. Plasmodb database (14) was then searched using their Plasmodb ID (PF3D7_1248000 and PF3D7_1454100 respectively), and putative proteins encoding for tRNA-splicing endonuclease (annotated as *Pf*TSEN1) and tRNA intron endonuclease (annotated as *Pf*TSEN2) were identified. tRNA-splicing endonuclease (*Pf*TSEN1) was studied for its structure-function characterization with a hypothesis that this enzyme may be responsible for cleaving the intron from pre-tRNA^Leu^ (Apicoplast). The PhenoPlasm database [18] indicated *Pf*TSEN1 essentiality for parasite survival, which underscores its importance in malaria parasite biology and makes it probable candidate for anti-malarial drug development. Sequence analysis and domain organization using Blastp (19) suggested that *P. falciparum* tRNA spicing endonuclease (*Pf*TSEN1) is 220 amino acids long and harbours a typical C-terminal catalytic domain (Fig. 2 A i). Domain organization of splicing endonucleases in other organisms including archaea, yeast and *Homo sapiens* also suggest the presence of a catalytic domain, responsible for the splicing endonuclease activity (Fig. 2 A ii, iii, iv). The presence of *Pf*TSEN1 homologues in other species was next investigated. Organism-specific Blastp search using *Pf*TSEN1 sequences provided their most reliable homologues in *archaea*, *Homo sapiens* and higher plants. Multiple sequence alignment of all sequences using Clustal omega (20) indicates that residues comprising the catalytic triad of *Pf*TSEN1 (His, Tyr and Lys) are fully conserved among all counterparts (Fig. S2). Motif analysis using MEME suit tool (21) further suggested the presence of different motifs involving the residues of catalytic triad (Fig. 2 B). Phylogenetic tree suggests that *Pf*TSEN1 and its homologues are grouped into different branches where *Pf*TSEN1 was observed to be more closely related to archael TSENs and distantly related to *Homo sapiens* (Fig. 2C). Gene location of *Pf*TSEN1 was identified to be on chromosome 12 (1973559 - 1975093 (+) (Fig. 2D i) while Gene location of pre-tRNA^(Leu)^ (Apicoplast) was found to be 809 - 1028 (+) (Fig. 2D ii).

**Figure 2:**
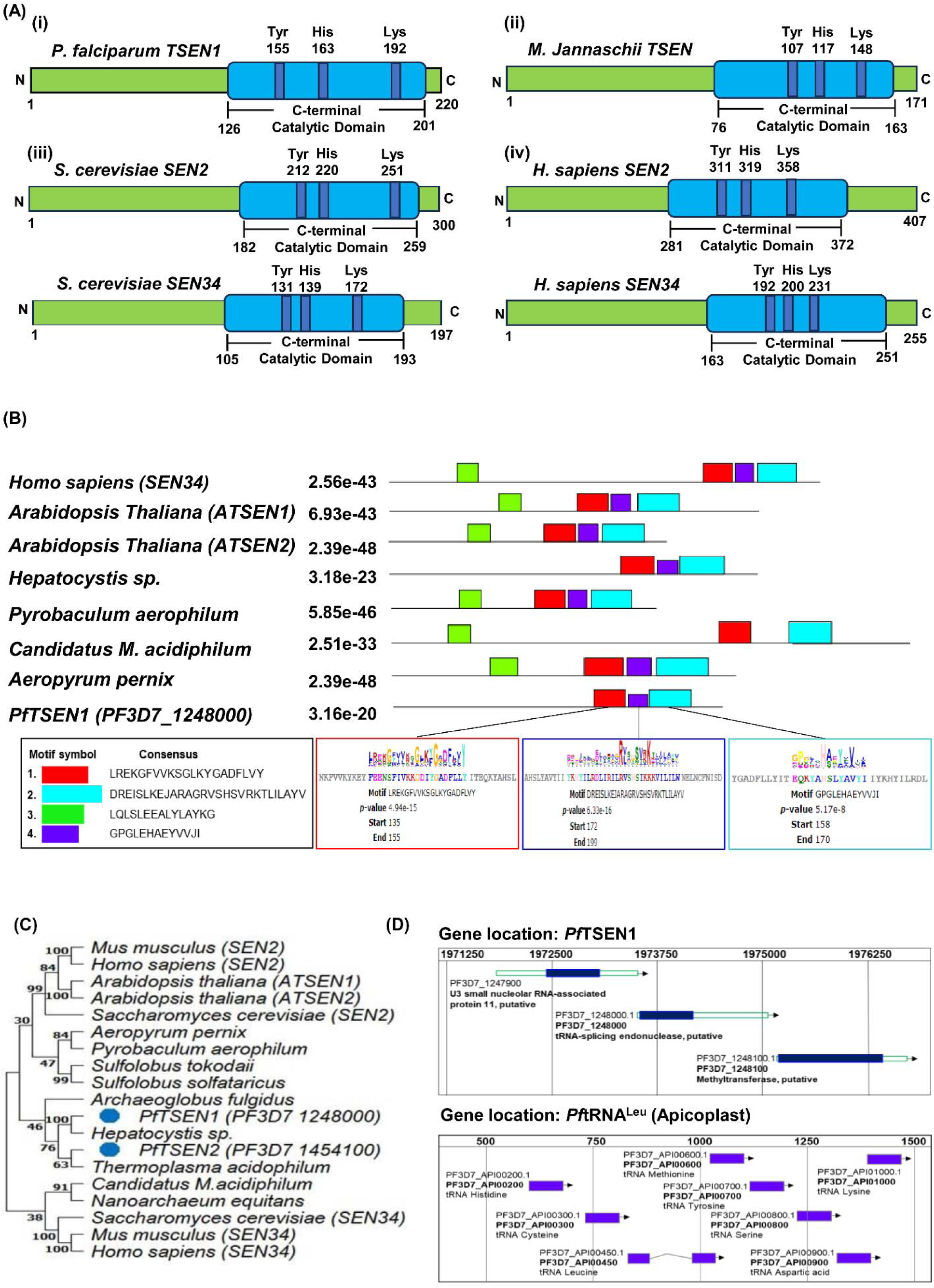
*In silico* analysis of *Pf*TSEN1. **(A)** Schematic representation of domain organization of *Pf*TSEN1 **(i)** and its homologs in archaea (*M. Jannaschii TSEN*) (**ii)**, yeast (*S. cerevisiae* SEN2, SEN34) **(iii)** and *H. sapiens (SEN2, SEN34)* **(iv)** depicting the C-terminal catalytic domain and the conserved catalytic triad of His, Lys, Tyr. **(B)** Motif prediction using MEME 5.5.4 and FIMO showing the presence of the consensus sequences within homologs of *Pf*TSEN1 in different species. **(C)** Phylogenetic tree demonstrating the evolutionary relationships of *Pf*TSEN1 with its homologs in other species using MEGA11. The evolutionary history was inferred using the Neighbor-Joining method and the evolutionary distances were computed using the Poisson correction method. **(D)** Schematic representation of genomic location of *Pf*TSEN1 (1973559 - 1975093 (+) and tRNA^Leu^ [Apicoplast; 809 - 1028 (+)].

### *Pf*TSEN1 shows co-expression at transcripts levels with tRNA substrate^Leu^ (apicoplast) at intra-erythrocytic stages and binds *in silico* to splice site of substrate

We next analysed the transcriptome data of *Pf*TSEN1 and pre-tRNA^Leu^ (Apicoplast) retrieved from Plasmodb database (14) that suggested their co-existence during asexual blood stages of malaria parasite (Fig. 3 A). Preliminary analysis of binding of *Pf*TSEN1 with pre-tRNA^Leu^ (Apicoplast) was performed by modelling the 3D structures of both enzyme and substrate. C-terminal region of *Pf*TSEN1 harboring the catalytic domain (120-201 aa position) was modelled using I-TASSER (Fig. 3 B) (22) whereas RNA structure modeling server ‘RNAComposer’ (23) was used to predict the 3D structure of pre-tRNA^Leu^ (Apicoplast) (Fig. 3 C). Enzyme-substrate docking studies performed using HDOCK (24) clearly demonstrated their interaction *in silico* (Fig. 3 D). Detailed residue analysis of docked complex suggested that catalytic triad residues (TYR155, HIS163 and LYS192) are directly involved in creating the interaction interface with substrate (Fig. 3 D). Tyr155 and His163 of *Pf*TSEN1 were observed to interact with A205, A206 of pre-tRNA^Leu^ (Apicoplast) whereas Lys192 of *Pf*TSEN1 binds with A204 of substrate. Interestingly, docking analysis further suggest that *Pf*TSEN1 residues (TYR166, LYS193, VAL194, ILE195, LEU196, ILE197) form receptor-ligand interface at nucleotide bases 174-176 of substrate which form nearby residues of 3’ splice site. 3D structure of mature tRNA^Leu^ (Apicoplast) without intron formed post splicing event is shown in fig. 3 E.

**Figure 3:**
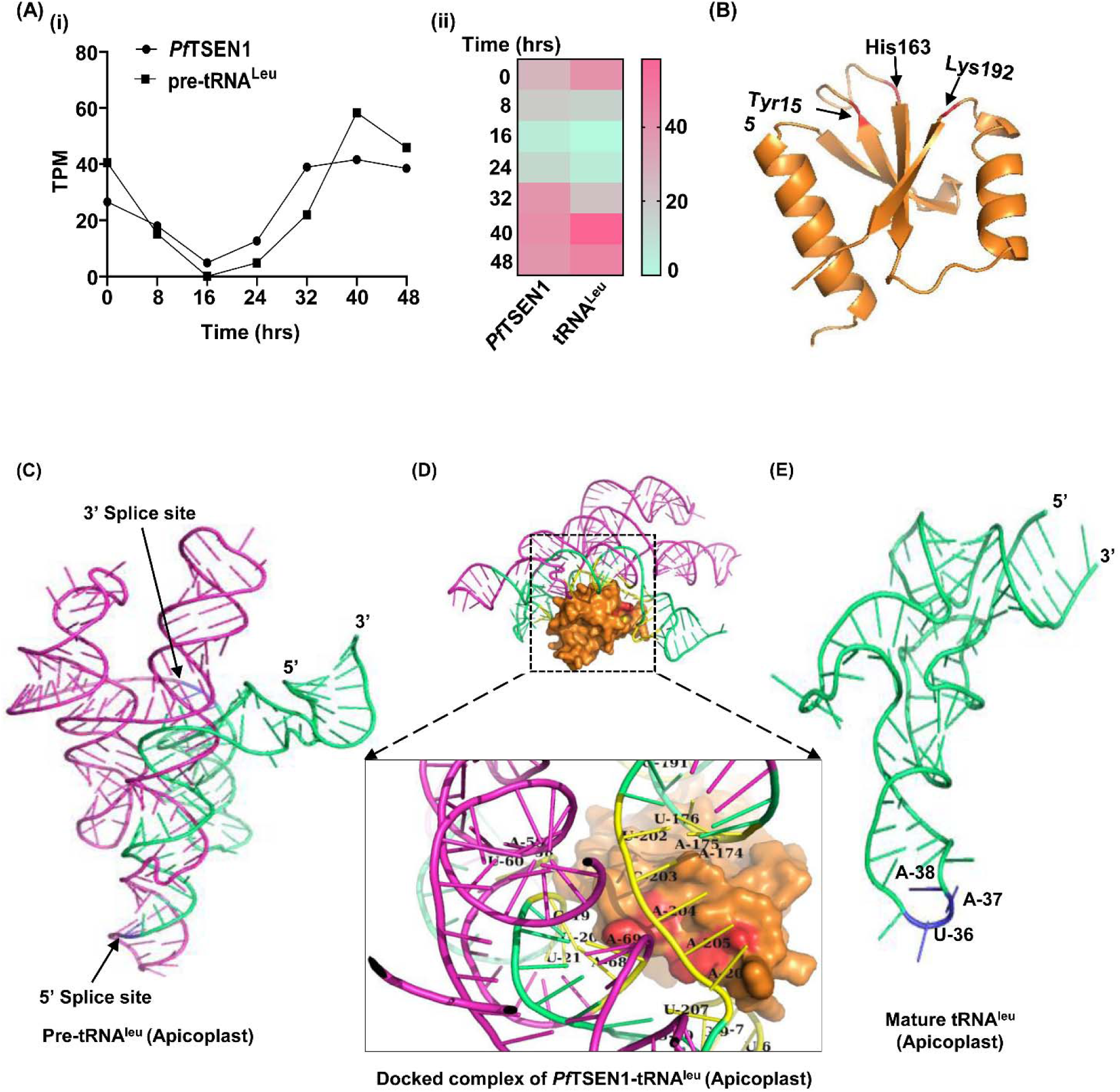
*In silico* binding of *Pf*TSEN1 with tRNA^Leu^ (Apicoplast). **(Ai)** Relative expression of *Pf*TSEN1 and tRNA^Leu^ (Apicoplast) at transcript levels during asexual blood stages of malaria parasite retrieved from Plasmodb database. **(ii)** Heat map depicting differential expression of *Pf*TSEN1 and tRNA^Leu^ (Apicoplast) at different intraerythrocytic stages. **(B)** Cartoon representation of homology model of *Pf*TSEN1 (120-201 aa position) predicted using I-TASSER. Residues forming the catalytic triad are mentioned on the model. **(C)** 3-dimenstional structural model of pre-tRNA^Leu^ (Apicoplast) showing the 5’ and 3’ splice sites. Exons (1-36, 172-220 nucleotides) are represented in green whereas intronic region (37-171 nucleotides) is depicted in magenta. **(D)** Docked complex of *Pf*TSEN1 with pre-tRNA^Leu^ (Apicoplast) showing the interaction interface. Surface representation of *Pf*TSEN1 is shown in orange while residues of catalytic triad are depicted in red. Zoom image of interaction interface is shown in box. **(E)** 3-dimenstional structural model of mature tRNA^Leu^ (Apicoplast) showing the anticodon formed (UAA) after splicing event.

### *Pf*TSEN1 is expressed in clinical stages of malaria and shows punctate expression

We next performed immunofluorescence assays to study the expression of *Pf*TSEN1 at asexual blood stages using anti-*Pf*TSEN1 antibodies. Thin blood smears of mixed stage *Pf*3D7 and cultures were fixed with methanol and blocked with 5% BSA in PBS. The slides were probed with anti-*Pf*TSEN1 antibodies (1:250) followed by Alexa Fluor 488 conjugated anti-mice secondary antibodies (1:500). The parasite nuclei were counterstained with DAPI (4′,6′-diamidino-2-phenylindole). Our preliminary fluorescence microscopy data suggested that *Pf*TSEN1 is expressed at ring, trophozoite and schizont stages of malaria parasite (Fig. 4 A i). We performed confocal microscopy at schizont stages to further study *Pf*TSEN1 expression and localization. The punctate fluorescence pattern near the DAPI stained nucleus observed for *Pf*TSEN1 was suggestive of its probable localization to apicoplast (Fig. 4 A ii). Furthermore, the protein sequence of *Pf*TSEN1 was analysed for the presence of apicoplast targeting leader sequence as reported by Foth et al (25). A presence of N terminal sequence of *Pf*TSEN1 aligning with apicoplast targeting leader sequence was observed. No staining was observed with pre-immune sera, suggesting the specificity of antibodies used in the assay. (Fig. 4 A iii). Expression analysis of *Pf*TSEN1 during the intra-erythrocytic parasite stages was also investigated using specific antisera against *Pf*TSEN1. Immunoblot analysis on total protein extracts of mixed stage *Pf*3D7 lysate showed a single protein band at ∼ 25 kDa for *Pf*TSEN1 (Fig. 4 B). No band was detected with pre-immune sera as depicted in fig. 4B ii.

**Figure 4:**
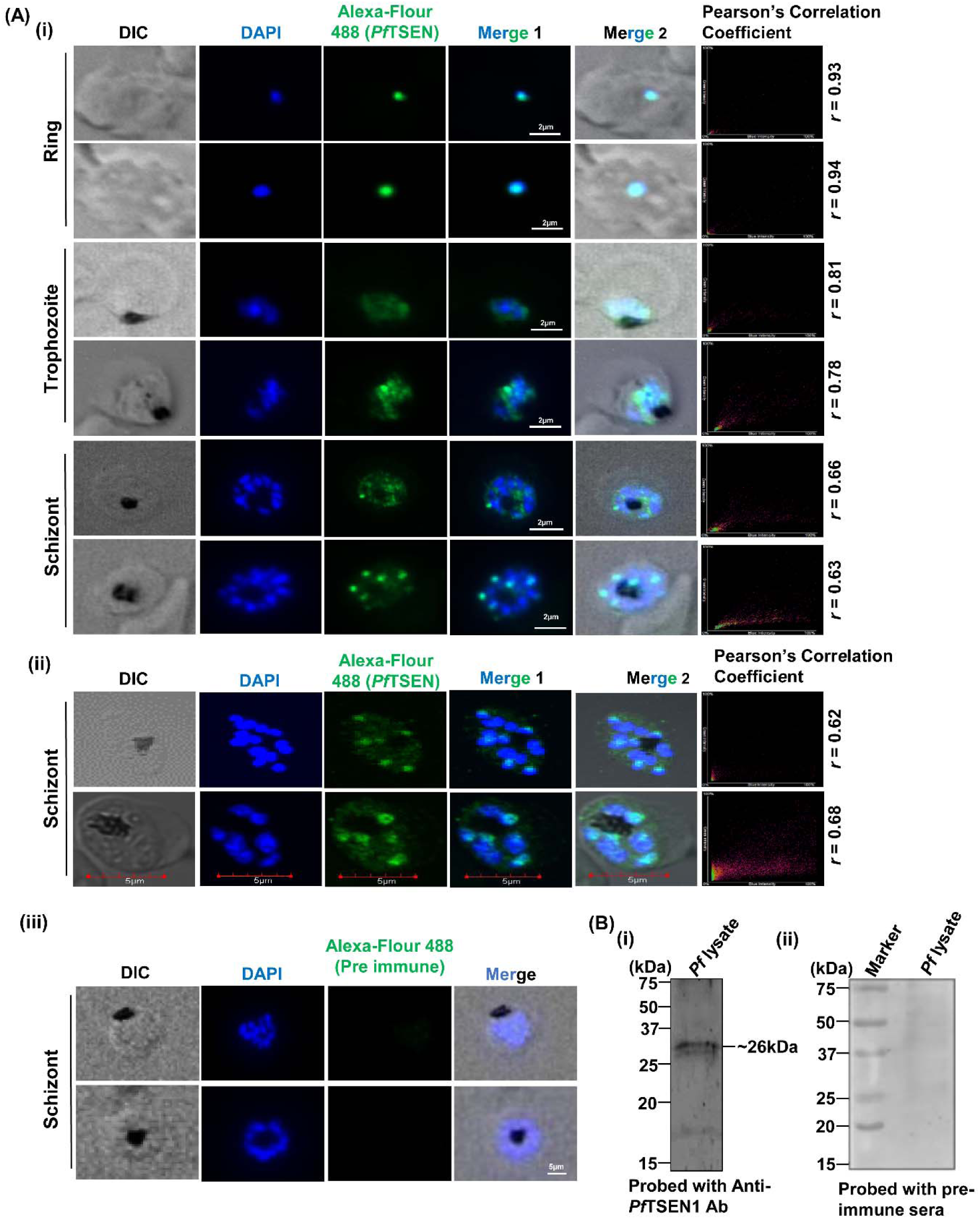
Expression and localization analysis of *Pf*TSEN1 at asexual blood stages of parasite life cycle. **(A) (i)** Fluorescence microscopy images showing the localization of *Pf*TSEN1 at asexual stages of parasite life cycle. Smears of methanol-fixed *Pf*3D7-infected erythrocytes were stained with anti-*Pf*TSEN1 antibodies (1:200) followed by incubation with Alexa Fluor-conjugated secondary antibodies (Alexa Fluor 488, green, dilution: 1:250). DIC: differential interference contrast image, DAPI with antifade: nuclear staining 40, 6-diamidino-2-phenylindole (blue); *Pf*TSEN: mouse anti-*Pf*TSEN1 (green); merge 1: overlay of *Pf*TSEN1 with DAPI (Scale bar: 2 µm). merge 2: overlay of DIC with *Pf*TSEN1 and DAPI (Scale bar: 2 µm). Plots representing Pearson’s correlation coefficient (r) of represented images are shown. **(ii)** Immunofluorescence images showing the localization of *Pf*TSEN1 at asexual stages of parasite life cycle by confocal microscopy. Smears of methanol-fixed *Pf*3D7-infected erythrocytes were stained with anti-*Pf*TSEN1 antibodies (1:250) followed by incubation with Alexa Fluor-conjugated secondary antibodies (Alexa Fluor 488, green, dilution: 1:250). DIC: differential interference contrast image, DAPI: nuclear staining 40, 6-diamidino-2-phenylindole (blue); *Pf*TSEN1: mouse anti-*Pf*TSEN (green); merge 1: overlay of *Pf*TSEN1 with DAPI (Scale bar: 5 µm), merge 2: overlay of DIC with *Pf*TSEN1 and DAPI (Scale bar: 2 µm). Plots representing Pearson’s correlation coefficient (r) of represented images are shown. **(iii)** Immunofluorescence images of schizonts probed with pre-immune sera as control. **(B i)** Immunoblot representing a band of expected molecular weight for *Pf*TSEN1 in parasite lysate. Sample was run on 15 % SDS-PAGE and subjected to Western blotting by probing with mice anti-*Pf*TSEN1 antibody. **(ii)** Western blot of parasite lysate probed with pre-immune sera as negative control.

### *Pf*TSEN1 exhibits splicing endonuclease activity to catalyse the formation of mature tRNA from pre-tRNA

Full length gene encoding for *Pf*TSEN1 was cloned in pMTSAra vector and recombinant protein was expressed in the soluble form in *E. coli* BL21 (DE3) cells with a 6X hexahistidine tag. Overexpression of the protein was induced with 0.4% L-Arabinose at 0.6 O.D. of bacterial secondary culture. Recombinant *Pf*TSEN1 was purified using Ni-NTA affinity chromatography (Fig. 5A i) and identity of protein was verified by western blotting using anti-histidine antibodies (Fig. 5 A ii). Purified *Pf*TSEN1 run as species of approximately 25kDa on SDS-PAGE (Fig. 4A i). Male BALB/c mice were used to produce polyclonal antisera against *Pf*TSEN1. A far-UV CD spectra on *Pf*TSEN1 was collected that depict the correctly folded state of recombinant protein (Fig. 5A iii).

**Figure 5.**
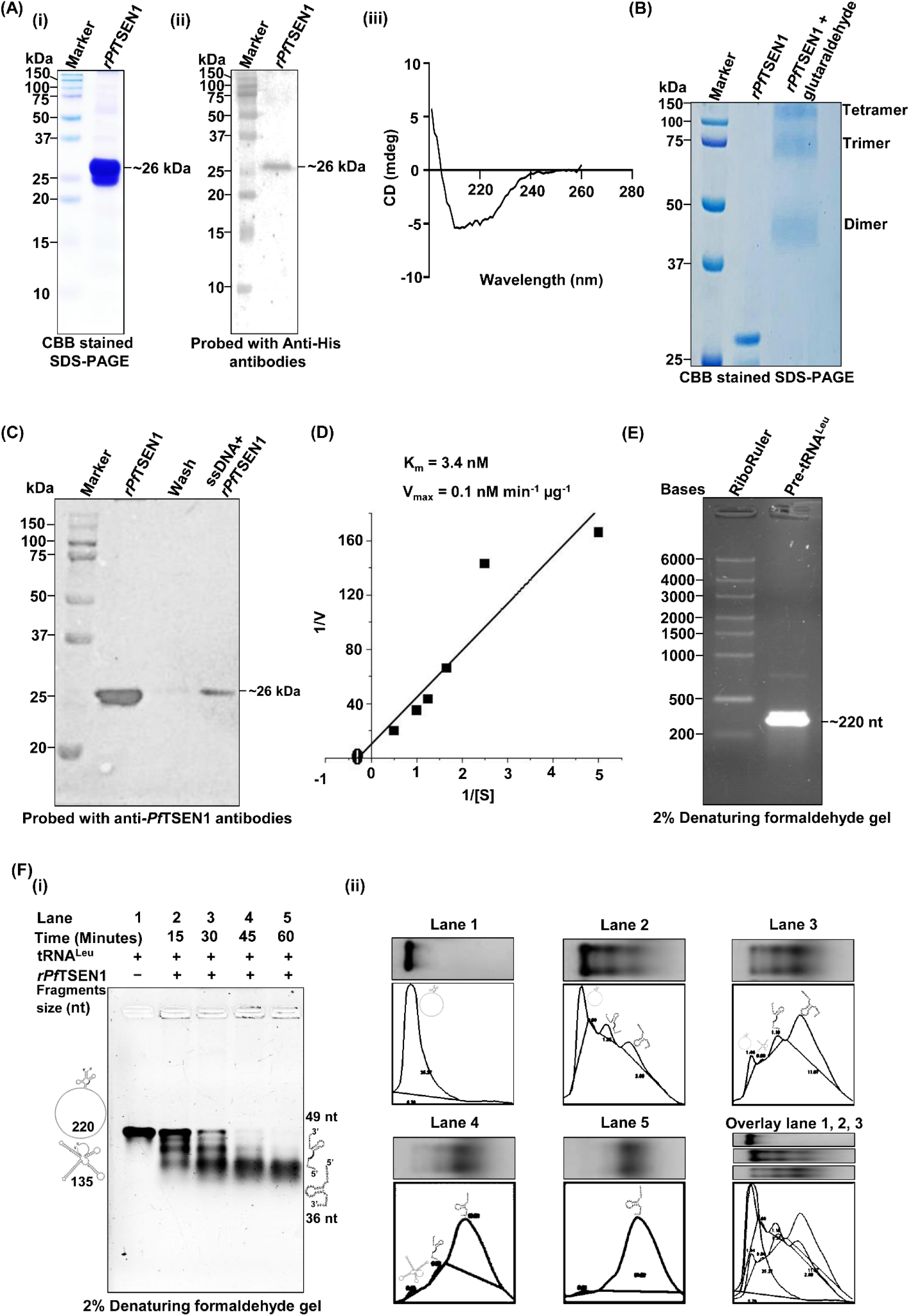
Expression and purification of *Pf*TSEN1 and its *in vitro* splicing endonuclease activity assays. **(A) (i)** SDS-PAGE showing recombinant purified *Pf*TSEN1 tagged with 6x-Histidine and stained with Coomassie Brilliant Blue (CBB). **(ii)** Western blot analysis of recombinant purified *Pf*TSEN1 probed with anti-histidine antibodies (1:5000). **(iii)** Secondary structure analysis of *Pf*TSEN1 using CD spectroscopy. The CD spectra of *Pf*TSEN1 was collected at 260 to 200 nm range and plotted with CD (mdeg) against its wavelength (nm). **(B)** SDS-PAGE showing oligomeric forms of *Pf*TSEN1 after incubation with 2.5% glutaraldehyde. Different conformations of protein are labelled. **(C)** *Pf*TSEN1-RNA interaction using pull down assay. *Pf*TSEN1 was incubated with single stranded (ss) DNA oligomer immobilized on cellulose beads in a pull down assay. Elutes and washes from the assay were loaded on 15% SDS-PAGE and subjected to western blotting using anti-*Pf*TSEN1 antibodies. **(D)** Phosphodiesterase assay of *Pf*TSEN1. Bis(p-nitrophenyl) phosphate was used as substrate (S) and incubated with *Pf*TSEN1 to determine the phosphodiesterase activity. Reaction velocity (v) was calculated at different substrate concentrations ([S]) and plotted according to the graphical representation of the Michaelis-Menten equation: 1/v = K_m_ /(V_max_ [S]) + 1/V max. A straight line was obtained using Origin software through linear curve fitting and subsequently, V max and K_m_ were calculated from the Y-intercept and the slope of the line respectively. **(E)** 2% denaturing formaldehyde gel depicting the *in vitro* synthesized and purified intron containing pre-tRNA^Leu^ (Apicoplast) substrate. **(F i)** 2% denaturing formaldehyde gel showing the cleavage of intron containing substrate pre*-Pf*tRNA^Leu^ (Apicoplast) by *Pf*TSEN1. Representative time courses (15, 30, 45, 60 minutes) for incubation of *Pf*TSEN1 with substrate are mentioned on the plot. **(ii)** Band intensity plots of pre-tRNA^Leu^ (Apicoplast) with and without *Pf*TSEN1 in splicing endonuclease assay using ImageJ. Lanes and gel images are shown above the intensity plots for reference.

We employed glutaraldehyde crosslinking assays to investigate the oligomeric confirmation of *Pf*TSEN1. Here, recombinant *Pf*TSEN1 was treated with 2.5% glutaraldehyde at 37°C for 30 mints and post treatment, sample was run on 12 % SDS-PAGE. On crosslinking, a band corresponding to dimeric and trimeric and tetrameric conformation of *Pf*TSEN were observed (Fig. 5B). Next, we used pull-down assays to test RNA binding properties of *Pf*TSEN1. Cellulose beads containing immobilized calf thymus ssDNA were mixed with 1 μg *Pf*TSEN1 and incubated at 4□ for 30 minutes. Bead pellet was washed thrice and boiled in 1X SDS-PAGE sample loading dye. The samples were loaded on SDS-PAGE and subjected to Western blotting and probed with anti-*Pf*TSEN1 antibodies. A band for *Pf*TSEN1 was seen in boiled beads sample, suggestive of its interaction with ssDNA (Fig. 5C). Biochemical assays were performed to test the phosphodiesterase activity of *Pf*TSEN1 using Bis(*p*-nitrophenyl) phosphate as substrate. The release of *p*-nitrophenol was monitored at 405 nm and reaction velocity (*v*) was calculated at different substrate concentrations (0.2 mM – 2 mM) and plotted according to the graphical representation of the Michaelis-Menten equation as described in methods section. K_m_ and V_max_ of *Pf*TSEN1 was found to 3.4nM and 0.1nM min^-1^ µg^-1^ be respectively (Fig. 5D).

Next we prepared pre-tRNA substrate *in vitro* using TranscriptAid T7 High Yield Transcription Kit, and analysed the splicing endonuclease activity of *Pf*TSEN1 using splicing assays. Transcribed tRNA^Leu^ (Apicoplast) substrate was purified using GeneJET TM RNA Purification Kit (Thermo Scientific TM, USA) (Fig. 5 E). Endonuclease cleavage activity of the recombinant *Pf*TSEN1 was determined by incubating purified recombinant protein with substrate at 37°C for time points of 15, 30, 45 and 60 minutes. The reactions were stopped by adding 2x RNA loading dye and samples were separated on 2% denaturing formaldehyde agarose gel. Our results suggested that upon incubating *Pf*TSEN1 with pre-tRNA^Leu^ substrate, distinct bands of cleaved products were seen depicting the ability of *Pf*TSEN1 to cleave intron-containing tRNA substrates (Fig. 5F i). Band intensities of pre-tRNA^Leu^ (Apicoplast) with and without *Pf*TSEN1 in splicing endonuclease assay were plotted using ImageJ that further demonstrate that the substrate is spliced out for intron removal (Fig. 5F ii).

### A novel synthetic compound ‘TSEN ^’^ inhibits tRNA slicing endonuclease activity of *Pf*TSEN1

In order to chemically target *Pf*TSENs and inhibit their splicing endonuclease activity, we carried out literature survey and found a unique class of compounds known as naphthoquinones that exhibit pleiotropic properties including antimicrobial, antitumor, antifungal, antiviral and antiprotozoal (26). A report by Karkare et al, suggested that naphtoquinones (*e.g.* diospyrin and 7-methyljuglone) bind to *M. tuberculosis* gyrase ‘GyrB’ at a novel site which is close to the ATPase site (27). Another study by Kennedy et. al, showed that naphthoquinone adduct 12,13-dihydro-N-methyl-6,11,13-trioxo-5H-benzo[4,5]cyclohepta[1,2-b] naphthalen-5,12-imine (TU100) possess a novel topoisomerase I/II inhibitory activity and exhibits chemotherapeutic potential (28). In light of these facts that naphthoquinones and their derivatives bind with DNA-binding proteins, we synthesized a novel derivative of naphtoquinone ‘1-(3-aminopropyl)-1H-pyrrolo[2,3-f]quinoline-4,9-dione’ (termed as TSEN_i_, Fig. 6A) to test its binding and inhibitory role on the functional activity of *Pf*TSEN1 considering its RNA binding properties. NMR spectra and synthesis of TSEN_i_ are represented in fig. S3, S4, S5.

**Figure 6:**
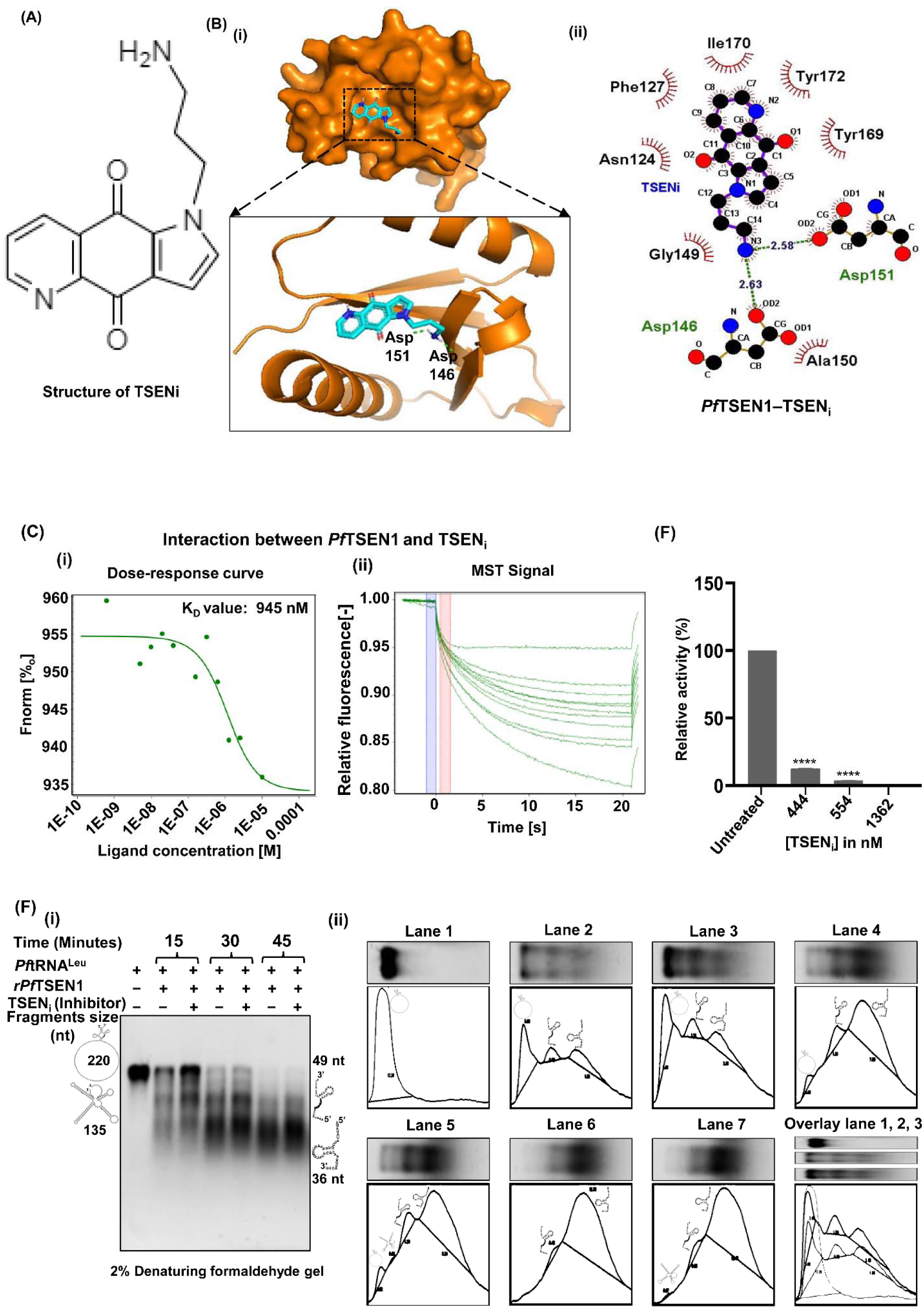
Interaction of TSEN_i_ with *Pf*TSEN1 and its inhibitory effect on splicing endonuclease activity of *Pf*TSEN1. **(Ai)** Schematic representation of chemical structure of TSEN_i_. **(ii)** Surface representation of *Pf*TSEN1-TSEN_i_ docked complex representing interaction interface of both protein and inhibitor. *Pf*TSEN1 is represented in orange colour and TSEN_i_ is represented in cyan colour. **(iii)** Ligplot showing the residues of *Pf*TSEN1 involved in interaction with TSEN_i_. **(C)** Interaction studies of *Pf*TSEN1 with TSENi using Microscale Thermophoresis. Labelled *Pf*TSEN1 was titrated against varying concentrations of TSEN_i_. Dose-response curve was generated that resulted in K_d_ value of 945 nM for *Pf*TSEN-TSENi interaction. **(D)** Inhibition of phosphodiesterase activity of *Pf*TSEN1 by TSEN_i_. *Pf*TSEN1 was incubated with different concentrations of TSEN_i_ (444, 545 and 1362 nM). Post incubation, Bis(p-nitrophenyl) phosphate was added as substrate and release of *p*-nitrophenol was monitored at 405 nm. Enzyme activity of *Pf*TSEN1 was considered 100% in absence of TSEN_i_. Relative activity of *Pf*TSEN1 in the presence of TSEN_i_ was calculated with respect to control with no inhibitor and plotted. **(E) (i)** Inhibition of splicing endonuclease activity of *Pf*TSEN1 by TSEN_i_. *Pf*TSEN1 was incubated with TSEN_i_ (1µM) before adding pre-tRNA^Leu^ (apicoplast). Representative time courses (15, 30, 45 minutes) are shown for the incubation of *Pf*TSEN1 with substrate. Samples were separated on 2% denaturing formaldehyde gel and are depicted. **(ii)** Band intensity plots of pre-tRNA^Leu^ (Apicoplast) in presence of TSEN_i_ in splicing endonuclease assay using ImageJ. Lanes and gel images are shown above the intensity plots for reference.

We performed preliminary analysis of binding by docking *Pf*TSEN1 with TSEN_i_ using AutoDock 4.2.6 (29). A stable *Pf*TSEN1-TSEN_i_ docked complex with a binding energy of -6.39 kcal/mol was observed (Fig. 6 Bi). Interface residues of *Pf*TSEN1 was observed within the active site region of *Pf*TSEN1 (145-192 aa). Asp146 and Asp151 of *Pf*TSEN1 were found to form H-bonds with TSEN_i_. Ligplot depicting the Hydrogen bonds and hydrophobic interactions between *Pf*TSEN1-TSEN_i_ docked complex is shown in fig. 6B ii. To validate *Pf*TSEN1 binding with TSEN_i_, we performed kinetic analysis of one-to-one interaction of purified recombinant *Pf*TSEN1 with TSEN_i_ using MST by NanoTemper Monolith NT.115 instrument. MST is a biophysical technique that relies on binding-induced changes in thermophoretic mobility and depends on different molecular properties such as particle size, charge, conformation, hydration state and solvation entropy. Upon titrating TSEN_i_ against fluorescently labelled recombinant *Pf*TSEN1, a gradual change in thermophoretic mobility was observed indicating interaction of TSEN_i_ with *Pf*TSEN1 (Fig. 6C). A K_d_ value of 945 nM was found, indicating good binding strength of TSEN_i_ with *Pf*TSEN1. Fig. 6 C shows dose response curves and MST signals for *Pf*TSEN1-TSEN_i_ binding.

We next analysed the effect of TSEN_i_ on the phosphodiesterase activity of *Pf*TSEN1. Here three different concentrations of TSEN_i_ [444nM (8xIC_50_), 545nM (10xIC_50_), 1362nM (25x IC_50_)] were incubated with *Pf*TSEN1 for 2 hrs. Post incubation, Bis(*p*-nitrophenyl) phosphate was added as substrate and release of *p*-nitrophenol was monitored at 405 nm. Relative enzyme activity of *Pf*TSEN1 in the presence of TSEN_i_ was calculated with respect to control with no inhibitor. Our data suggest that enzyme activity of *Pf*TSEN1 is greatly reduced on incubation with TSEN_i_, suggesting the inhibitory potential of TSEN_i_ on the activity of *Pf*TSEN1 (Fig. 6D). Also, the decline in the activity of *Pf*TSEN1 was enhanced on increasing the TSEN_i_ concentration. We also performed tRNA intron splicing assay of *Pf*TSEN1 in presence and absence of TSEN_i_ to test the inhibitory effect of TSEN_i_ on splicing endonuclease activity of the enzyme. Here, recombinant *Pf*TSEN1 (5 μg) was incubated with TSEN_i_ (1 μM) for 2 hrs and subsequently with substrate pre-tRNA^Leu^ (Apicoplast; 5 μg) at 37°C for the various time points (15, 30 and 45 minutes). The reactions were stopped by adding 2x RNA loading dye and samples were separated on 2% denaturing formaldehyde agarose gel. Our results suggested that upon adding TSEN_i_, the band intensity of pre-tRNA^Leu^ substrate was higher as compared to untreated samples at different time points, indicating the inability of *Pf*TSEN1 to cleave the substrate (Fig. 6E i). These data demonstrate that TSEN_i_ has potent inhibitory effect on the splicing endonuclease activity of *Pf*TSEN1. Band intensities of pre-tRNA^Leu^ (Apicoplast) in the presence and absence of TSEN_i_ in splicing endonuclease assay were plotted using ImageJ that further demonstrate that *Pf*TSEN1 splicing endonuclease activity is inhibited via TSEN_i_ (Fig. 6F ii).

### Blocking splicing endonuclease activity of *Pf*TSEN1 by TSEN_i_ inhibits parasite growth

TSEN_i_ was screened for its anti-plasmodial activity against the asexual stage of the human malaria parasite *in vitro*. Highly synchronized ring-stage parasites of 3D7 and Dd2 were exposed to a range of compound concentration (250 nM-1 µM) for 72 hrs. Parasitaemia was measured at 72 hrs post-treatment using SYBR Green I based fluorescence assay and the graph was plotted for the average value of three independent sets of experiments. Schematic representation of the growth inhibition assay in depicted in fig. 7 A i. Our data revealed the potent anti-malarial activity of TSEN_i_ against 3D7 and Dd2 strain of *P. falciparum* with a significant reduction in the parasite load (Fig. 7 A ii, iii). IC_50_ value of TSEN_i_ in 3D7 and Dd2 strain of *P. falciparum* was found to be 54.54 nM and 49.20 nM respectively. Giemsa stained images of *Pf*3D7 and Dd2 treated with and without TSEN_i_ are shown in fig. 7 A iv, v.

**Figure 7.**
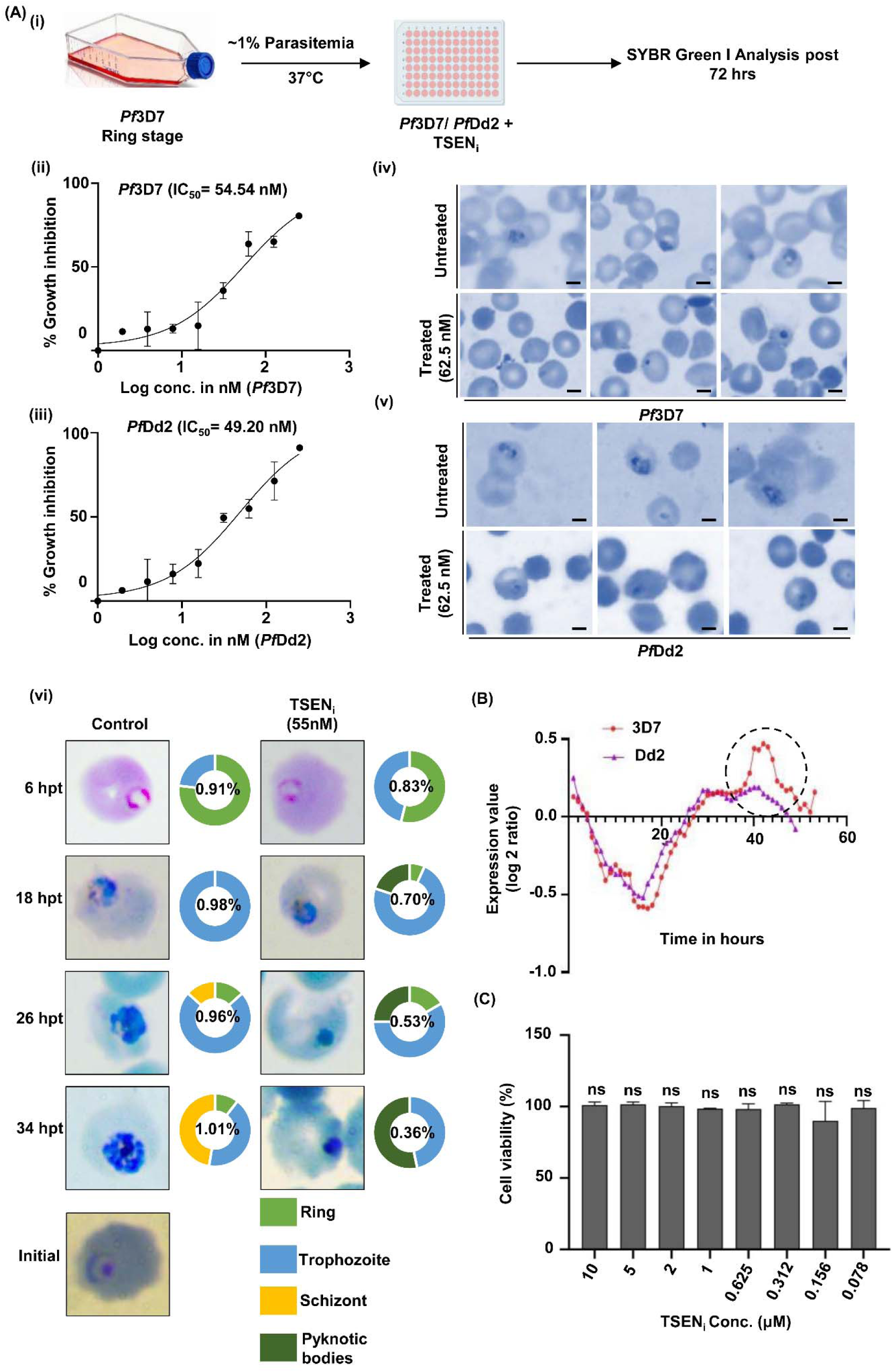
Growth inhibitory effect of TSEN_i_ on *Pf*3D7 and *Pf*Dd2. **(A i)** Schematic representation of methodology used in the assay. The parasite culture synchronized at ring stage was treated with different concentrations (250, 125, 62.5, 31.25, 15.62, 7.8, 3.9, 1.95, 1.00 nM) of TSEN_i_ for 72 hrs, and the percent growth inhibition was estimated. **(ii)** IC_50_ value of TSEN_i_ in *Pf*3D7 were evaluated by plotting growth inhibition values against the log concentration of TSEN_i_. The experiment was done in triplicate, and the results were shown as mean values ± SD. IC_50_ value is mentioned on the plot. **(iii)** IC_50_ value of TSEN_i_ in *Pf*Dd2 were evaluated by plotting growth inhibition values against the log concentration of TSEN_i_. The experiment was done in triplicate, and the results were shown as mean values ± SD. IC_50_ value is mentioned on the plot. **(iv, v)** Giemsa-stained images of *Pf*3D7 and *Pf*Dd2 treated with and without TSEN_i_. **(vi)** Growth progression assays of TSEN_i_. *Pf*3D7 ring staged parasites at ∼1% parasitemia and 2% hematocrit were treated with 55nM of TSEN_i_. Blood smears were drawn at different time points post treatment (hpt), fixed with methanol, and stained with giemsa solution. The progression of parasite was monitored by manually counting the parasitaemia at each stage. Percentage parasitaemia of controls and treated cultures are mentioned on the plot. Pie charts represents the ratio of asexual blood stages of parasite at different time points post TSEN_i_ treatment. **(B)** Transcriptomics data of *Pf*TSEN1 in *Pf*3D7 and chloroquine resistant strain *Pf*Dd2 during intra-erythrocytic stages retrieved from Plasmodb database. **(C)** MTT cell-viability assay of TSEN_i_ on HepG2 cells. Cell viability of untreated cells were considered 100%. Relative cell viability of treated cells were calculated and plotted as bar graph.

We next investigated stage specific effect of TSEN_i_ on *Pf*3D7 using growth progression assays. *Pf*3D7 ring staged parasites at ∼1% parasitemia was treated with TSEN_i_ at their IC_50_ concentration and blood smears were drawn at different time points post treatment, fixed with methanol, and stained with giemsa solution. Untreated culture was used as a negative control. Growth progression defect was analysed by counting ∼2000 cells per giemsa stained slide. In untreated culture, growth of the cells progressed normally whereas significant reduction of parasitemia at all three asexual stages of malaria parasite was found in TSEN_i_ treated culture as compared to untreated control (Fig. 7A v). Also, TSEN_i_ was found to effect the trophozoite stages of parasite and inhibit their progression to schizont stages.

We also analysed the transcriptomics data of *Pf*TSEN1 in chloroquine sensitive 3D7 and chloroquine resistant Dd2 strains of *Plasmodium* retrieved from Plasmodb database (14). The data depict similar expression profile of *Pf*TSEN1 in both strains (Fig. 7B). However, around 40 hrs post invasion of intra-erythrocytic cycle, *Pf*TSEN1’s expression was comparatively high in *Pf*3D7 as compared to Dd2 strain. These data further marks the importance of *Pf*TSEN1 in drug resistant strains and hints towards using this enzyme as a target for drug resistance management. Next, the toxicity of TSEN_i_ was investigated in the human hepatocellular carcinoma cell line, HepG2 by using the MTT cell-viability assay. At 10 μM concentration, TSEN_i_ displayed negligible toxicity on HepG2 cells upon 48 hrs of treatment, with treated cells as healthy as control ones (Fig. 7C).

## Discussion

RNA interactome in all living organisms form an important part of cell biology and cover all aspects of functions associated with RNAs including splicing, export, stability, translation and degradation (4–6). Assemblies of RNA and protein molecules form ribonucleoprotein (RNPs) complexes that mainly comprise RNA interactome, and play role in cellular adaptation. These RNPs contain RNAs ranging from messenger RNAs (mRNAs) to non-coding RNAs, rRNAs and tRNAs (4–6). In line with this, we attempted to identify the presence of total tRNAs in malaria parasite and the enzymes responsible for their splicing and maturation. Our data mining in *Plasmodium* identified 45 tRNAs in nucleus and 33 tRNAs in apicoplast. A previous report by Preiser et. al reported 25 tRNA genes transcribed from the plastid-like DNA (apicoplast) of *Plasmodium falciparum* (30). Our study has now provided the updated list of total tRNAs present in nucleus and apicoplast of malaria parasite. tRNA specific for leucine in apicoplast was observed to harbor 135 bases long intron (37 -171). The presence of intron containing tRNA highlighted the existence of tRNA splicing endonucleases responsible for intron splicing. A recent report by Hollin et al. performed a proteome-wide approach called R-DeeP, a method based on sucrose density gradient ultracentrifugation to identify RNA dependent proteins (RDPs) (4). We observed tRNA splicing endonucleases to be a part of these ribonucleoprotein (RNPs) complexes, and whose functional role in *Plasmodium* was totally undefined. Splicing endonucleases (TSENs) are important players in tRNA maturation in all living organisms since tRNA molecules are essentially required for translating genetic information into protein (9). A plethora of processing steps are required for the generation of functional tRNAs that involve removal of 5’ leader and 3’ trailer sequences from pre-tRNA. Additionally some pre-tRNAs contain introns that need to be removed to generate functionally mature tRNAs. These processing steps are performed by a splicing endonuclease that marks the importance of this enzyme in cell biology. Archaeal and eukaryotic TSENs are well characterized with their precise functions known (9). Both systems harbor a related TSEN, however the subunit composition has diverged. Based on the complex architecture, three types of TSENs are found in archaea *viz*. homotetrameric: α_4_, homodimeric: α_2_, and heterotetrameric: (αβ)_2_], each possess different substrate specificity during the splicing process (31). On the contrary, humans possess a single type of TSEN composed of four different subunits (αβγδ) (32). Based on current literature, archaea and eukaryotic TSENs are functionally known proteins with their defined roles well illustrated, however the existence and structure-function relevance of TSENs in *Plasmodium* is fully unexplored. Since TSENs are essential proteins and considered as probable druggable targets, our study for the first time attempted to elucidate the identification of TSENs in *Plasmodium falciparum* (*Pf*TSEN1 and *Pf*TSEN2) and provide evidence for structure-function studies of these novel molecules with special emphasis to *Pf*TSEN1.

Our search in Plasmodb database (14) identified two tRNA splicing endonucleases (*Pf*TSEN1, *Pf*TSEN2) in *Plasmodium* that harbor a C terminal catalytic domain with a conserved catalytic triad of Tyr, His, and Lys residues, accountable for cleaving the phosphodiester backbone. Motif analysis along with phylogenetic studies suggest that *Pf*TSEN1 is homologous to archael TSENs as compared to humans. The study was then sought to analyse the expression and localization of *Pf*TSEN1 at asexual blood stages of malaria parasite. Our immunofluorescence data clearly depict the expression of *Pf*TSEN1 during asexual blood stages of malaria parasite. The punctate fluorescence pattern observed near DAPI staining for *Pf*TSEN1 suggests its probable localization to apicoplast. Western blotting using anti-*Pf*TSEN1 antibodies depict the expression of *Pf*TSEN1 in parasite lysate. Further the expression of pre-tRNA^Leu^ (Apicoplast) substrate at schizont stages of *Pf*3D7 was confirmed by using RT-PCR.

We next tested the potential of *Pf*TSEN1 to exhibit phosphodiesterase and splicing endonuclease activity by using *in vitro* assays. Phosphodiestearse activity of *Pf*TSEN1 was tested by using Bis(*p*-nitrophenyl) phosphate as substrate and release of *p*-nitrophenol (yellow colored product) was monitored at 405 nm. Our data clearly suggested the ability of *Pf*TSEN1 to exhibit phosphodiesterase activity with a K and V of 3.4 nM and 0.1nM min^-1^ µg^-1^ respectively. V_max_ of *Pf*TSEN1 was identified to be low owing to the fact that Bis(*p*-nitrophenyl) phosphate is a small molecule that can diffuse from the catalytic cavity of *Pf*TSEN1. Splicing endonuclease activity of *Pf*TSEN1 was assessed by incubating purified recombinant protein with pre-tRNA^Leu^ (Apicoplast) substrate at 37°C for different time points. Post incubation, the samples were subjected to separate on 2% denaturing formaldehyde agarose gel. Presence of four distinct bands of substrate post incubation with protein was observed that clearly demonstrate the splicing endonuclease activity of *Pf*TSEN1.

Since *Pf*TSENs are probable druggable target and malaria elimination demands new anti-malarial drug development, we next attempted to search for specific inhibitor of *Pf*TSEN1 that can block its endonuclease activity function, and can ultimately obstruct the protein translation. Our literature search along with *in silico* docking analysis suggested a RNA binding protein inhibitor, which is a Napthoqinone derivative (annotated as ‘TSEN_i_’) to bind with *Pf*TSEN1. Interaction of TSEN_i_ with *Pf*TSEN1 was further confirmed by using MST which depicted a K_d_ value of 945 nM for TSEN_i_ - *Pf*TSEN1 interaction. We then assessed the inhibitory effect of TSEN_i_ on the splicing endonuclease activity of *Pf*TSEN1 by incubating the protein with the inhibitor prior to its incubation with pre-tRNA^Leu^ substrate. Our data depict that TSEN_i_ can block the splicing endonuclease function of *Pf*TSEN1 as the band intensity of substrate was higher in the presence of TSEN_i_ as compared to its absence. These data prompted us to test the anti-plasmodial activity of TSEN_i_ on asexual blood stages of malaria parasite. Growth inhibition assays illustrated that TSEN_i_ inhibited the growth of clinical stage *P. falciparum* 3D7 and chloroquine drug-resistant strain Dd2 with an IC_50_ value of 54.54 nM and 49.20 nM respectively. Also, TSEN_i_ was observed to effect the trophozoite stages of parasite and inhibit their progression to schizont stages.

Overall, our study provide first evidence for the existence of tRNAs and tRNA splicing endonucleases in *Plasmodium falciparum* that function to excise introns from pre-tRNAs to yield mature tRNAs. Based on our data and proposed functions of mature tRNAs, we have proposed a model depicting the role of *Pf*TSEN1 in parasite during the clinical stages of development (Fig. 8). Since formation of mature tRNAs are necessity of cells to pursue protein translation, identification and characterization of splicing endonucleases responsible for tRNA maturation in *Plasmodium falciparum* forms a key step towards understanding malaria biology, and provide a druggable target for therapeutics development. In line with this, our work has identified a naphthoquinone derivative as an inhibitor of *Pf*TSEN1 that showed potent anti-malarial profile, and can possibly be combined with known anti-malarials without losing its efficacy.

**Fig. 8.**
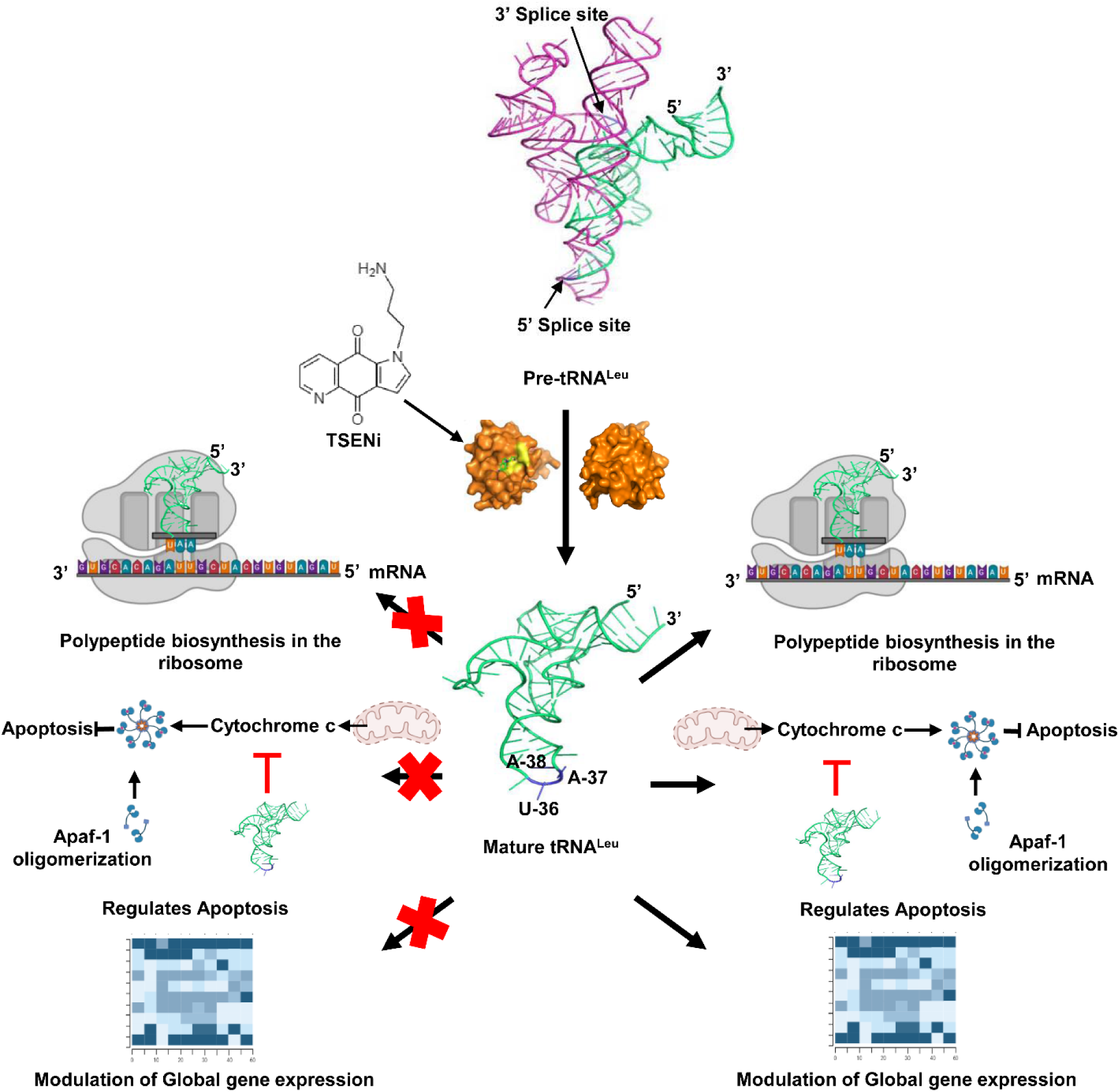
Model diagram depicting the functional role of *Pf*TSEN1 in malaria parasite and its inhibition by TSEN_i_ blocks the formation and functions of mature tRNA.

## Methodology

### In silico analysis of Plasmodium tRNAs

The presence of total tRNAs in *Plasmodium falciparum 3D7* was searched using Plasmodb database (14). The secondary structures of pre-tRNAs were predicted using RNAfold webserver (16) and were visualized and developed using webserver R2DT (https://rnacentral.org/r2dt) (33) and RNAcanvas (https://rnacanvas.app) (34) respectively. The 2D structures were further converted and stabilized into 3D structures using RNA structure modeling server, RNAComposer (<u>rnacomposer.ibch.poznan.pl</u>) (23, 35, 36).

### Real-time PCR analysis of tRNA^Leu^ (Apicoplast and nuclear)

Expression of tRNA^Leu^ (Apicoplast) and tRNA^Leu^ (nuclear) at transcript levels were evaluated during asexual blood stages of *Pf*3D7 using Real-Time PCR (StepOnePlusTM Real-Time PCR system, Applied Biosystems, USA). 18S rRNA was taken as a positive control. The primer sequences for the real-time PCR analysis of substrates and 18S rRNA are mentioned in Table S3. The reaction mixture (10 μl) comprised of cDNA, 5 μl SYBR™ Green PCR Master Mix (Applied Biosystems™), and 1 μl (5 μM) tRNA^Leu^ (Apicoplast) specific forward and reverse primers. The PCR conditions included initial denaturation at 95°C for 5 min, followed by amplification for 40 cycles of 15 seconds each at 95°C, 5 seconds at 55°C, and 1 minute at 72°C, with fluorescence acquisition at the end of each extension step. Amplification was immediately followed by a melt program consisting of 15 seconds at 95°C, 1 minute at 60°C, and a stepwise temperature increase of 0.3°C/s until 95°C, with fluorescence acquisition at each temperature transition. All samples were evaluated in duplicates.

### *In silico* analysis of *Pf*TSEN1

Protein sequences of *Pf*TSEN1 (PF3D7_1248000) was retrieved from PlasmoDB database (https://Plasmodb.org) (14). Protein sequence similarity search was performed with BLASTp tool using non-redundant database search (19). The conserved domains were identified using CDD search tool of NCBI (37, 38) and motifs were predicted using MEME v 5.5.5 (Multiple Em for Motif Elicitation) (21) and FIMO v 5.5.5 (Find Individual Motif Occurrence) (39). Multiple sequence alignment between archaeal TSENs and eukaryotes TSENs were performed using Clustal Omega (20). Evolutionary relationships were analysed using Neighbor-joining method in MEGA11 and iTOL (interactive Tree of Life, https://itol.embl.de) (40, 41).

### *In silico* modeling of *Pf*TSEN1 and its docking with pre-tRNA^Leu^ (apicoplast)

3D model of C-terminal catalytic domain of *Pf*TSEN1 was considered due to the functional importance of evolutionary conserved catalytic triad in the tRNA splicing endonuclease(s). The 3D structure of C-terminal catalytic region (120-201 aa position) of *Pf*TSEN1 was modelled using I-TASSER (22). This 3D model of 82 amino acids was further submitted to ModRefiner for protein structure refinement (42). The stereochemical quality of this 3D structure was analyzed for its overall residue by residue geometry using PROCHECK server (43). Protein-tRNA molecular interaction studies between C-terminal *Pf*TSEN1 and pre-tRNA^Leu^ (apicoplast) was performed using HDOCK (http://huanglab.phys.hust.edu.cn/) (24). HDOCK is based on a hybrid algorithm and performs *ab initio* free docking using template-based modeling. Docked complexes were visualized using PyMOL (44).

### Immunofluorescence assay (IFA)

IFAs were performed on synchronized *Plasmodium falciparum* 3D7 culture to check the expression and localization of *Pf*TSEN1. Thin blood smears of mixed asexual stage *Pf*3D7 cultures (8-10% parasitemia) were fixed with pre-chilled methanol for overnight at -20°C. Non-specific binding sites in the parasite were blocked with 3% BSA (in 1×PBS), for 1 hr at room temperature and probed with polyclonal anti-*Pf*TSEN1 mouse serum at dilutions (1:250) for 1 hr at room temperature (RT). Slides were washed twice with 1xPBS containing 0.05% Tween-20 (PBST), followed by washing once with 1xPBS, and probed with Alexa-Fluor 488 conjugated anti-mouse IgG (Molecular Probes, U.S.A.; 1:250) for 1 hr at RT. All antibodies were diluted in 1% BSA, prepared in 1X PBST. The slides were washed and mounted with Fluoroshield antifade reagent with DAPI (40,6-diamidino-2-phenylindole; Sigma). Images were acquired in Olympus FLUOVIEW FV3000 confocal microscope using Olympus cellSens dimensions imaging software.

### *In vitro* expression analysis of *Pf*TSEN1 in parasite

*Pf*3D7 mixed stage asexual cultures with ∼8-10% parasitemia were harvested and subjected to saponin lysis (0.15% w/v) followed by extensive washing of parasite pellet with 1x PBS to remove haemoglobin. Purified parasites were subjected to RIPA lysis using two volume of RIPA lysis buffer (30 mM Tris, 150 mM NaCl, 20 mM MgCl_2_, 1mM EDTA, 1 mM β-ME, 0.5% Triton X-100, 1% IGEPAL, 1 mM PMSF, and 0.1% SDS; pH 8.0). Sample was separated by SDS-PAGE and analyzed by Western blot analysis by probing with mouse anti-*Pf*TSEN1 polyclonal antibody (1:250), followed by HRP-conjugated secondary anti-mice antibody (1:5000). Blot was developed by using ECL substrate.

### Cloning, expression and purification of *Pf*TSEN1

Full length *Pf*TSEN1 and *Pf*TSEN2 was codon-optimized and synthesized in pUC57 through GenScript, with the gene cloned at the unique SnaBI restriction site. Both gene inserts were excised from pUC57 and cloned in SnaBI-cut dephosphorylated pMTSAra plasmid. The positive clones were selected by colony PCR and confirmed by plasmid restriction digestion and nucleotide sequencing. The cloned plasmids were transformed in BL21 *E. coli* strain and overexpression of *Pf*TSEN1 was induced with 0.4% L-Arabinose at 0.6 OD of bacterial secondary culture, for 18 hrs at 16°C. Protein purification was performed using Ni-NTA affinity chromatography.

### Raising polyclonal antisera against *Pf*TSEN1

To raise specific antibodies against *Pf*TSEN1 protein, three female Balb/c mice (6 to 8 weeks old) were administered (i.p.) with 12 μg of the recombinant protein (in 0.9% saline), in a prime and boost regimen. For the priming dose (day 0), the formulation was made by thoroughly mixing equal volumes of Freund’s complete adjuvant and saline containing the protein. For subsequent booster doses (days 21 and 42), formulations were made with Freund’s incomplete adjuvant. Blood samples were collected from the retro-orbital sinus of the mice on days 31 and 52 (terminal bleed) after primary immunization. Collected blood samples were incubated for 30 min at 37°C, centrifuged at 1200×g at 4°C for 15 min, and then serum samples were collected and stored at −80°C until further analysis.

### Circular dichroism (CD) spectroscopy

The far-UV CD spectra were collected on *Pf*TSEN1 by chiarascan using 1 mm cuvette. Protein was diluted to 0.2mg/ml concentration in phosphate buffer, pH 7.4 and the CD data was collected in 260 to 200 nm wavelength range with 1 nm bandwidth and 40 nm/min scanning rate. Five scans were collected and averaged. The baseline was subtracted from the averaged CD data. The CD (mdeg) was plotted against wavelength.

### Glutaraldehyde cross-linking assays

Recombinant *Pf*TSEN1 (∼30 µg) were suspended in crosslinking buffer (50 mM sodium phosphate pH 8, 100 mM NaCl) in a total volume of 50 µl. 5µl of 2.3% glutaraldehyde was added to the reaction and incubated at 37°C for 30 minutes. The reaction was terminated by adding 10µl of stop solution (1 M Tris-HCl, pH 8.0), and samples were resolved on 10% SDS-PAGE.

### Bead based *Pf*TSEN1-RNA pull down assay

To determine whether *Pf*TSEN1 binds to RNA, we performed a pull-down assay using cellulose beads with immobilized single stranded calf thymus DNA (Sigma Aldrich, USA). 5 μg recombinant *Pf*TSEN1 was incubated with 500 μl cellulose matrix (1mg/ml; dissolved in 50 mM NaH_2_PO_4_ (pH 7.2), 150 mM NaCl buffer) and incubated at 4□ for 1 hr. The bead pellet was washed extensively with 250µl buffer (50 mM NaH_2_PO_4_, pH 7.2 and 150 mM NaCl). Post washing, beads were boiled in 1X SDS-PAGE sample loading dye and loaded on 15% SDS-PAGE. Samples were subjected to Western blotting and probed with anti-*Pf*TSEN1 antibodies followed by HRP-conjugated secondary anti-mice antibody (1:5000). Blot was developed by using ECL substrate.

### Phosphodiesterase activity assays of *Pf*TSEN1 in presence and absence of TSEN_i_

Bis(p-nitrophenyl) phosphate was utilized as the substrate to determine the phosphodiesterase activity. A typical 500 µl reaction mixture contains 20 mM Tris-HCl (pH 7.4), substrate of indicated concentration (0.2 mM – 2 mM), 13 µg protein, and 0.2 mM ZnCl_2_. The release of p-nitrophenol (□ = 11500 M^−1^ cm^−1^ at pH 7.4) was continuously monitored at 405 nm. One unit of activity corresponds to 1 μmol of p-nitrophenol liberated per minute at 37 °C. Reaction velocity (v) was calculated at different substrate concentrations ([S]) and plotted according to the graphical representation of the Michaelis-Menten equation: 1/v = K_m_ /(V_max_ [S]) + 1/V max. A straight line was obtained using Origin software (43) through linear curve fitting and subsequently, V max and K_m_ were calculated from the Y-intercept and the slope of the line respectively.

The effect of TSEN_i_ on the phosphodiesterase activity of *Pf*TSEN1 was analysed by incubating different concentrations of TSEN_i_ [444nM (8xIC_50_), 545nM (10xIC_50_), 1362nM (25x IC_50_)] with *Pf*TSEN1 (13 µg) for 2 hrs at 37°C. Post incubation, Bis(*p*-nitrophenyl) phosphate was added as substrate (2mM) and release of *p*-nitrophenol was monitored at 405 nm. As a control, *Pf*TSEN1 was added to substrate without TSEN_i_ and its activity was considered 100%. Relative activity of *Pf*TSEN1 in the presence of TSEN_i_ was calculated with respect to control with no inhibitor and plotted as bar graph.

### Preparation of synthetic tRNA substrate

Pre-tDNA^Leu^ (Apicoplast) sequence was amplified from genomic DNA isolated from the *Plasmodium falciparum* (3D7) using specific primer pairs (Table S1). The Amplified products were excised and purified from the gel using Thermo Scientific GeneJET Plasmid Miniprep Kit (Thermo Scientific). tRNA^Leu^ (Apicoplast) sequence was transcribed from the respective tDNA template using TranscriptAid T7 High Yield Transcription Kit (Thermo Scientific, USA) and the transcribed RNA was purified through GeneJET RNA Purification Kit (Thermo Scientific, USA).

### tRNA intron splicing assay

The cleavage of pre-tRNA^Leu^ (Apicoplast) was carried out in a 20 μl reaction mixture, containing 5 μg of purified recombinant *Pf*TSEN1, 5 μg of substrate, and 1x reaction buffer (25 mM Tris-HCl, pH 8, 20 mM KCl, 6 mM MgCl_2_, BSA 5 mg/ml, 0.5% glycerol, 1 mM DTT, 0.5 mM EDTA). The reaction mixture was incubated at a temperature of 37°C for time points of 15, 30, 45 and 60 minutes. The reactions were stopped by adding 2x RNA loading dye (95% formamide, 0.025% SDS, 0.025% bromophenol blue, 0.025% xylene cyanol FF, 0.025% ethidium bromide, 0.5mM EDTA) and was heated at 70°C for 10 minutes. The samples were separated on 2% denaturing formaldehyde agarose gel (prepared in 10X MOPS buffer with a composition of 0.4 M MOPS (pH 7.0), 0.1 M Sodium acetate, 0.01 M EDTA (pH 8.0) and 37% formaldehyde solution) and visualized under UV light using a ChemiDoc Imaging System (Bio-Rad). Intensity plots of bands obtained were prepared using ImageJ (46).

### *In silico* docking of *Pf*TSEN1 with TSEN_i_

*In silico* docking of *Pf*TSEN1 with TSEN_i_ was performed with Lamarckian Genetic Algorithm (LGA) using AutoDock 4.2.6 (28). Residue interaction interface of docked complex was analysed using Ligplot (47). Docked complexes were visualized using PyMOL (43).

### Binding assays using Microscale Thermophoresis (MST)

To determine the binding between *Pf*TSEN1 and TSEN_i_, MST analysis was performed by using Monolith NT.115 instrument (NanoTemper Technologies, Munich, Germany). Briefly, 15 μM *Pf*TSEN1 was labelled using 30 μM Lysine reactive dye following standard protocol (NanoTemper’s Protein Labelling Kit RED-NHS 2^nd^ Generation), and incubated in dark for 30 minutes at RT. Fractions of labelled protein were collected and subjected to fluorescent counts. Interacting ligand TSEN_i_ (20 μM) was serially diluted in decreasing concentration, and titrated in HEPES-NaCl/0.05% tween-20 in a 1:1 ratio against the constant concentration of *Pf*TSEN1 (375 nM). Pre-mixed sample was incubated at RT for 15 minutes, followed by centrifugation at 10,000 rpm for 10 minutes. The samples were loaded into the standard capillaries (K002 Monolith NT.115) and thermophoretic mobility was measured. Data evaluation was performed with the Monolith software (Nano Temper, Munich, Germany).

### Effect of TSEN_i_ on *Pf*TSEN1 endonuclease activity

To test the inhibitory effect of TSEN_i_ on endonuclease activity of *Pf*TSEN1, tRNA slicing assay was performed in the presence and absence of TSEN_i._ Here 1 μM TSEN_i_ was first incubated with 5 μg of recombinant *Pf*TSEN1 for 30 mints on ice. Subsequently, 5 μg of substrate pre-tRNA^Leu^ (Apicoplast) in 1x reaction buffer (25 mM Tris-HCl, pH 8, 20 mM KCl, 6 mM MgCl2, BSA 5 mg/ml, 0.5% glycrol, 1 mM DTT, 0.5 Mm EDTA) was added and incubated at 37°C for various time points (15, 30 and 45 mints). The reactions were stopped by adding 2x RNA loading dye (Thermo Scientific^TM^, USA) and was heated at 70°C for 10 minutes. The Samples were separated on 2% denaturing formaldehyde agarose gel and visualization under UV light using ChemiDoc Imaging System (Bio-Rad).

### Growth inhibition assay

The anti-plasmodial activity of TSEN_i_ was tested on 3D7 and Dd2 strain of *Plasmodium falciparum*. Sorbitol synchronized ring stage cultures with 0.8-1% parasitemia and 2% hematocrit were dispensed into the plates and incubated for 72 hrs in a final volume of 100 µL/well. Different concentrations (250, 125, 62.5, 31.25, 15.62, 7.8, 3.9, 1.95 nM) of TSEN_i_ were added and post incubation parasite growth was determined using SYBR Green I based fluorescence assay. Here culture was lysed by freeze-thaw followed by addition of 100 µL lysis buffer (20 mM Tris/HCl, 5mM EDTA, 0.16% (w/v) saponin, 1.6% (v/v) Triton X) containing 1× SYBR Green I. Plates were incubated in dark at room temperature (RT) for 3-4 hrs. *Plasmodium falciparum* proliferation was assessed by measuring the fluorescence using a Varioskan™ LUX multimode microplate reader (Thermo Scientific™) with an excitation and emission of 485 nm & 530 nm respectively. IC_50_ values were determined via non-linear regression analysis using GraphPad Prism 8.0 software. The results were expressed as the percent inhibition compared to the untreated controls and calculated using the following formula:

100× ([OD of Untreated sample – blank]-[OD– blank]/ [OD – blank]). As blank, uninfected RBCs were used.

### Growth Progression Assay

To study the effect of TSEN_i_ on progression of parasites, *Pf*3D7 ring staged parasites at ∼1% parasitemia and 2% hematocrit were treated with 55nM of TSEN_i_ in a 96 well microtiter plate. Untreated parasite culture was kept as control and the assay plate was kept at 37°C under mixed gas condition, to enable parasite growth. Blood smears were drawn at different time points post treatment, fixed with methanol, and stained with giemsa solution. The progression of parasite was monitored by manually counting the parasitemia at each stage. A total of 2000 red blood cells were counted to estimate the stage specific parasitemia.

### MTT assay

Cytotoxic effect of TSEN_i_ on HepG2 was assessed by utilizing the ability of live cells to cleave MTT [(3-[4,5-dimethylthiazol-2-yl]-2,5-diphenyl tetrazolium bromide)] into blue-colored formazan crystals. Briefly, HepG2 cells were seeded in a 96-well microtiter plate at a density of 10,000 cells per 100 μL per well. Cells were allowed to proliferate at 37°C for 48 hrs in the presence of TSEN_i_ (10 μM). Cells cultured in the absence of any compound were maintained as the positive control. After treatment, MTT labelling reagent was added to each well to a final concentration of 0.5 mg/ml and incubated at 37°C for 4 hrs. Purple-colored formazan crystals, thus formed, were dissolved in 100 ul DMSO solvent and optical density was measured spectrophotometrically by taking absorbance at 570 nm using Varioskan™ LUX multimode microplate reader (Thermo Fisher Scientific™). The absorbance of untreated cells was taken as 100%.

## Acknowledgements

MKM is ICMR-SRF and **AB** is supported by DBT-Research Associateship (DBT-RA/2023/ January/N/3456). **AKK** is CSIR SRA in scientist pool scheme. Advance instrumentation and research facility at Jawaharlal Nehru University, New Delhi is acknowledged for providing confocal microscopy and Surface plasmon resonance facility.

## Author contributions

**MKM, AB, AKK:** Experimental design, experimentation, data analysis and manuscript writing. **FDL:** Synthesis of TSEN_i_. **PM and RN:** performed phosphodiesterase assays. **PJ:** experimentation and manuscript preparation. **JS:** performed immunofluorescence assays. **GK:** conducted growth inhibition assays. **AM:** conducted growth progression assays**. RS:** Performed MST experiments. **NJ, JP:** Preparation of buffers and manuscript preparation. **TD:** Data analysis and manuscript writing. **CA:** Syntheses of TSEN_i_, Data analysis and manuscript writing. **AR and SS:** Conception of idea, experimental design, data analysis and manuscript writing.

## Data availability statement

Any additional information supporting the current study in this paper is available from the corresponding author upon request.

## References

1. Venkatesan, P. (2024) The 2023 WHO World malaria report. Lancet Microbe. 10.1016/s2666-5247(24)00016-8.

2. Fairhurst, R.M. and Dondorp, A.M., (2016). Artemisinin-resistant Plasmodium falciparum malaria. Microbiol. Spectr. 4, 10.1128/microbiolspec. EI10-0013-2016.

3. Talisuna, A. O., Karema, C., Ogutu, B., Juma, E., Logedi, J., Nyandigisi, A., Mulenga, M., Mbacham, W. F., Roper, C., Guerin, P. J., D’Alessandro, U., and Snow, R. W. (2012) Mitigating the threat of artemisinin resistance in Africa: Improvement of drug-resistance surveillance and response systems. Lancet Infect Dis. 10.1016/S1473-3099(12)70241-4

4. Hollin, T., Abel, S., Banks, C., Hristov, B., Prudhomme, J., Hales, K., Florens, L., Stafford Noble, W. and Le Roch, K.G., 2024. Proteome-Wide Identification of RNA-dependent proteins and an emerging role for RNAs in Plasmodium falciparum protein complexes. Nature Communications, 15(1), p.1365.

5. Bleichert, F. and Baserga, S.J., 2010. Ribonucleoprotein multimers and their functions. Critical reviews in biochemistry and molecular biology, 45(5), pp.331–350.

6. Briata, P. and Gherzi, R., 2020. Long non-coding RNA-ribonucleoprotein networks in the post-transcriptional control of gene expression. Non-coding RNA, 6(3), p.40.

7. Geuens, T., Bouhy, D. and Timmerman, V., 2016. The hnRNP family: insights into their role in health and disease. Human genetics, 135, pp.851–867.

8. Calvin, K. and Li, H., 2008. RNA-splicing endonuclease structure and function. Cellular and Molecular Life Sciences, 65, pp.1176–1185.

9. Hayne, C. K., Sekulovski, S., Hurtig, J. E., Stanley, R. E., Trowitzsch, S., and van Hoof, A. (2023) New insights into RNA processing by the eukaryotic tRNA splicing endonuclease. Journal of Biological Chemistry. 10.1016/j.jbc.2023.105138

10. Xue, S., Calvin, K. and Li, H., 2006. RNA recognition and cleavage by a splicing endonuclease. Science, 312(5775), pp.906–910.

11. Zhang, X., Yang, F., Zhan, X., Bian, T., Xing, Z., Lu, Y. and Shi, Y., 2023. Structural basis of pre-tRNA intron removal by human tRNA splicing endonuclease. Molecular Cell, 83(8), pp.1328–1339.

12. Sekulovski, S., Sušac, L., Stelzl, L.S., Tampé, R. and Trowitzsch, S., 2023. Structural basis of substrate recognition by human tRNA splicing endonuclease TSEN. Nature Structural & Molecular Biology, 30(6), pp.834–840.

13. Avcilar-Kucukgoze, I., and Kashina, A. (2020) Hijacking tRNAs From Translation: Regulatory Functions of tRNAs in Mammalian Cell Physiology. Front Mol Biosci. 10.3389/fmolb.2020.610617

14. Aurrecoechea, C., Brestelli, J., Brunk, B.P., Dommer, J., Fischer, S., Gajria, B., Gao, X., Gingle, A., Grant, G., Harb, O.S. and Heiges, M., 2009. PlasmoDB: a functional genomic database for malaria parasites. Nucleic acids research, 37(suppl_1), pp.D539–D543.

15. Chan, P.P., Lin, B.Y., Mak, A.J. and Lowe, T.M., 2021. tRNAscan-SE 2.0: improved detection and functional classification of transfer RNA genes. Nucleic acids research, 49(16), pp.9077–9096.

16. Hofacker, I.L., 2003. Vienna RNA secondary structure server. Nucleic acids research, 31(13), pp.3429–3431.

17. UniProt Consortium. UniProt: a worldwide hub of protein knowledge. Nucleic acids research. 2019 Jan 8;47(D1):D506–15.

18. Sanderson, T. and Rayner, J.C., 2017. PhenoPlasm: a database of disruption phenotypes for malaria parasite genes. Wellcome open research, 2.

19. Altschul SF, Madden TL, Schäffer AA, Zhang J, Zhang Z, Miller W, Lipman DJ. Gapped BLAST and PSI-BLAST: a new generation of protein database search programs Nucleic Acids Res 25: 3389–3402.

20. Sievers F, Wilm A, Dineen D, Gibson TJ, Karplus K, Li W, Lopez R, McWilliam H, Remmert M, Söding J, Thompson JD. Fast, scalable generation of high□quality protein multiple sequence alignments using Clustal Omega. Molecular systems biology. 2011 Jan 1;7(1).

21. Bailey TL, Elkan C. Fitting a mixture model by expectation maximization to discover motifs in bipolymers.

22. Yang, J., Yan, R., Roy, A., Xu, D., Poisson, J. and Zhang, Y., 2015. The I-TASSER Suite: protein structure and function prediction. Nature methods, 12(1), pp.7–8.

23. Purzycka, K.J., Popenda, M., Szachniuk, M., Antczak, M., Lukasiak, P., Blazewicz, J. and Adamiak, R.W., 2015. Automated 3D RNA structure prediction using the RNAComposer method for riboswitches1. In Methods in enzymology (Vol. 553, pp. 3–34). Academic Press.

24. Yan Y, Zhang D, Zhou P, Li B, Huang SY. HDOCK: a web server for protein-protein and protein-DNA/RNA docking based on a hybrid strategy. Nucleic Acids Res. 2017 Jul 3;45(W1):W365–W373. doi: 10.1093/nar/gkx407. PMID: 28521030; PMCID: PMC5793843.

25. Foth, B.J., Ralph, S.A., Tonkin, C.J., Struck, N.S., Fraunholz, M., Roos, D.S., Cowman, A.F. and McFadden, G.I., 2003. Dissecting apicoplast targeting in the malaria parasite *Plasmodium falciparum*. Science, 299(5607), pp.705–708.

26. Babula, P., Adam, V., Havel, L. and Kizek, R., 2007. Naphthoquinones and their pharmacological properties. Ceska a Slovenska farmacie: casopis Ceske farmaceuticke spolecnosti a Slovenske farmaceuticke spolecnosti, 56(3), pp.114–120.

27. Karkare, S., Chung, T.T., Collin, F., Mitchenall, L.A., McKay, A.R., Greive, S.J., Meyer, J.J., Lall, N. and Maxwell, A., 2013. The naphthoquinone diospyrin is an inhibitor of DNA gyrase with a novel mechanism of action. Journal of Biological Chemistry, 288(7), pp.5149–5156.

28. Kennedy, S., DiCesare, J.C. and Sheaff, R.J., 2011. Topoisomerase I/II inhibition by a novel naphthoquinone containing a modified anthracycline ring system. Biochemical and biophysical research communications, 408(1), pp.94–98.

29. Eberhardt, J., Santos-Martins, D., Tillack, A.F. and Forli, S., 2021. AutoDock Vina 1.2. 0: New docking methods, expanded force field, and python bindings. Journal of chemical information and modeling, 61(8), pp.3891–3898.

30. Preiser, P., Williamson, D.H. and Wilson, R.J.M., 1995. tRNA genes transcribed from the plastid-like DNA of Plasmodium falciparum. Nucleic acids research, 23(21), pp.4329–4336.

31. Fujishima, K., Sugahara, J., Miller, C.S., Baker, B.J., Di Giulio, M., Takesue, K., Sato, A., Tomita, M., Banfield, J.F. and Kanai, A., 2011. A novel three-unit tRNA splicing endonuclease found in ultrasmall Archaea possesses broad substrate specificity. Nucleic acids research, 39(22), pp.9695–9704.

32. Hayne, C.K., Schmidt, C.A., Haque, M.I., Matera, A.G. and Stanley, R.E., 2020. Reconstitution of the human tRNA splicing endonuclease complex: insight into the regulation of pre-tRNA cleavage. Nucleic acids research, 48(14), pp.7609–7622.

33. Sweeney, B.A., Hoksza, D., Nawrocki, E.P., Ribas, C.E., Madeira, F., Cannone, J.J., Gutell, R., Maddala, A., Meade, C.D., Williams, L.D. and Petrov, A.S., 2021. R2DT is a framework for predicting and visualising RNA secondary structure using templates. Nature Communications, 12(1), p.3494.

34. Johnson, P.Z. and Simon, A.E., 2023. RNAcanvas: interactive drawing and exploration of nucleic acid structures. Nucleic acids research, 51(W1), pp.W501–W508.

35. Biesiada, M., Purzycka, K.J., Szachniuk, M., Blazewicz, J. and Adamiak, R.W., 2016. Automated RNA 3D structure prediction with RNAComposer. RNA Structure Determination: Methods and Protocols, pp.199–215.

36. Sarzynska, J., Popenda, M., Antczak, M. and Szachniuk, M., 2023. RNA tertiary structure prediction using RNAComposer in CASP15. *Proteins: Structure*, Function, and Bioinformatics, 91(12), pp.1790–1799.

37. Wang, J., Chitsaz, F., Derbyshire, M.K., Gonzales, N.R., Gwadz, M., Lu, S., Marchler, G.H., Song, J.S., Thanki, N., Yamashita, R.A. and Yang, M., 2023. The conserved domain database in 2023. Nucleic Acids Research, 51(D1), pp.D384–D388.

38. Marchler-Bauer, A., Bo, Y., Han, L., He, J., Lanczycki, C.J., Lu, S., Chitsaz, F., Derbyshire, M.K., Geer, R.C., Gonzales, N.R. and Gwadz, M., 2017. CDD/SPARCLE: functional classification of proteins via subfamily domain architectures. Nucleic acids research, 45(D1), pp.D200–D203.

39. Grant CE, Bailey TL, Noble WS. FIMO: scanning for occurrences of a given motif. Bioinformatics. 2011 Apr 1;27(7):1017–8.

40. Saitou, N. and Nei, M., 1987. The neighbor-joining method: a new method for reconstructing phylogenetic trees. Molecular biology and evolution, 4(4), pp.406–425.

41. Tamura, K., Stecher, G. and Kumar, S., 2021. MEGA11: molecular evolutionary genetics analysis version 11. Molecular biology and evolution, 38(7), pp.3022–3027.

42. Xu, D. and Zhang, Y., 2011. Improving the physical realism and structural accuracy of protein models by a two-step atomic-level energy minimization. Biophysical journal, 101(10), pp.2525–2534.

43. Laskowski, R.A., MacArthur, M.W., Moss, D.S. and Thornton, J.M., 1993. PROCHECK: a program to check the stereochemical quality of protein structures. Journal of applied crystallography, 26(2), pp.283–291.

44. DeLano, W.L., 2002. Pymol: An open-source molecular graphics tool. CCP4 Newsl. Protein Crystallogr, 40(1), pp.82–92.

45. Edwards, P.M., 2002. Origin 7.0: scientific graphing and data analysis software. Journal of chemical information and computer sciences, 42(5), pp.1270–1271.

46. Abràmoff, M.D., Magalhães, P.J. and Ram, S.J., 2004. Image processing with ImageJ. Biophotonics international, 11(7), pp.36–42.

47. Wallace, A.C., Laskowski, R.A. and Thornton, J.M., 1995. LIGPLOT: a program to generate schematic diagrams of protein-ligand interactions. Protein engineering, design and selection, 8(2), pp.127–134.

