## supplementary data for "A first-in-*Plasmodium* study on tRNA intron splicing endonuclease ‘*Pf*TSEN1’ and its substrate expression in clinical stage malaria"

*Corresponding information

**Table S1: List of tRNAs present in apicoplast of *Plasmodium falciparum 3D7*. Red color depicts the intronic region**

| S.No. | PlasmoDB ID | tRNA type | Anticodon loop (5'-3') | Exon | Transcript length | Location | Genomic Sequence |
|---|---|---|---|---|---|---|---|
| 1. | PF3D7_API00450 | Leucine | TAA | 2 | 85 | 809 - 1028 (+) | AGAGATATGGTGAAATTTGGTATACACAATGGACTTaaaataatttgagttaattattattaatattaaattttaagaaaatatataatatattttttaaattctgtaatatattttaaaatttatatattcaaaagactttatttataaaaagtctaaatttattaagAAAATCCATTAACATTATTGTTGTAAGGGTTCAAATCCCTTTATCTCTA |
| 2. | PF3D7_API05400 | Leucine | TAG | 1 | 79 | 27114 - 27192 (+) | ATGAATATGGCGAAATAGGTAAACGCACTAAATTTAGATTTTAGTTATTATAATAAGAGTTCAAATCTCTTTATTCATA |
| 3. | PF3D7_API06200 | Leucine | TAG | 1 | 79 | 30999 - 31077 (-) | ATGAATATGGCGAAATAGGTAAACGCACTAAATTTAGATTTTAGTTATTATAATAAGAGTTCAAATCTCTTTATTCATA |
| 4. | PF3D7_API00200 | Histidine | GTG | 1 | 72 | 643 - 714 (+) | ATAAATATAATCTAATGGTTAAGATGAAGAATTGTGGTTTCTTTTATATGAGTTCAAATCTCTTTATTTATC |
| 5. | PF3D7_API00300 | Cysteine | GCA | 1 | 71 | 733 - 803 (+) | AATGATATAACTTAATTGATAAAGTAAATAATTGCAAATTATTATATTTCAGTTTGAATCTGAATATCATT |
| 6. | PF3D7_API00600 | Methionine | CAT | 1 | 87 | 1034 - 1120 (+) | AACATTTATAGCTAAGTGGTCGAAAGCAATGGACTCATAATTCATTTTCATATATTGATCATCAGTAGTTCGAATCTACTTAAATGT |
| 7. | PF3D7_API00700 | Tyrosine | GTA | 1 | 84 | 1126 - 1209 (+) | AAAGTTAATGCCTGAGTGGTTAAAAGGAATGGACTGTAAATCCATTGATAATATATCTACATCAGTTCAAATCTGATTTAACTT |
| 8. | PF3D7_API00800 | Serine | GCT | 1 | 92 | 1223 - 1314 (+) | AAGAGAAATGACTGAGAGGTTTATAGTTATAAATTGCTAATTTATTGTATATATAATAATATTATACCAAGGGTTCGAATCCCTTTTTCTCT |
| 9. | PF3D7_API00900 | Aspartic acid | GTC | 1 | 76 | 1321 - 1396 (+) | AGAATTTGTAGTTTAATTAGGTAAAAATATTATTTTGTCATAATAAAGAATACGAGTTCAATTCTCGTCAAATTCG |
| 10. | PF3D7_API01000 | Lysine | TTT | 1 | 73 | 1407 - 1479 (+) | GAATTACTAGCTTAATTGGTAGAGTACTCGACTTTTAATCGAATGGTTCTGAGTTCAAATCTCAGGTAGTTCA |
| 11. | PF3D7_API01100 | Glutamic acid | TTC | 1 | 70 | 1487 - 1556 (+) | ACTTTTATCGTTTAAAGGTAAGACATCTTTTTTTCAAGAAGAAAATAGGAATTCGATTTTCCTTAAAAGT |
| 12. | PF3D7_API01200 | Proline | TGG | 1 | 72 | 1569 - 1640 (+) | CAGAATATAGTGTAATGGTAACATATCTATTTTGGGAATAGAAGAATATAGGTTCAAATCCTATTTTTCTGA |
| 13. | PF3D7_API03200 | Glutamine | TTG | 1 | 72 | 9967 - 10038 (+) | TAGAATATAACCAAAAGGTTAAGGTAATGAATTTTGATTTCATTAATATAGGTTCGAATCCTATTATTCTAA |

| | | | | | | | |
|---|---|---|---|---|---|---|---|
| **14.** | PF3D7_API03300 | Glycine | ACC | 1 | 70 | 10044 - 10113 (+) | ATGAATATAATTTAATGATAAAATACAATTTTACCATAATTGTT ATAAGAGTTTGAATCTCTTTATTCAT |
| **15.** | PF3D7_API03400 | Tryptophan | CCA | 1 | 71 | 10127 - 10197 (+) | ATGTCTTTAATTTAAAGTAAAAATATAAATTTCCAAAATTTATA ATAAAGGTTCGAATCCTTTAGGACATG |
| **16.** | PF3D7_API03700 | Glycine | TCC | 1 | 72 | 12906 - 12977 (+) | AACAAATATAGTTTAATCGGTAAAATATTAATTTTCCAAATTAA TGATATGGATTCAATTTCCATTATTTGT |
| **17.** | PF3D7_API03900 | Serine | TGA | 1 | 87 | 13246 - 13332 (+) | AAGAGAAATGACAGAGTGGTTTATTGTGTTTGATTTGAGATCAA AAAAATATAAATATATTTCATGGGTTCAAATCCCATTTTCTCT |
| **18.** | PF3D7_API04800 | Threonine | TGT | 1 | 73 | 23966 - 24038 (-) | GCTAAAGTAGCTTAATTGGTAAAGCAACTGATTTGTAATCAGTA GATTATGAGTTCAAATCTCACCATTAGCT |
| **19.** | PF3D7_API05000 | Methionine | CAT | 1 | 74 | 26791 - 26864 (-) | AGCGAAATAGAGCATAAGGAAAGTTCGTCGGATTCATGCTCCGA AGGTAATCGGTTCAATTCCGTTTTTCGCTT |
| **20.** | PF3D7_API05100 | Arginine | ACG | 1 | 72 | 26870 - 26941 (-) | AAACTTGTAATTTAATGGATAAAATATATAAATACGAATTATAA AATAAAAGTTCAATTCTTTTCAAGTTTA |
| **21.** | PF3D7_API05200 | Valine | TAC | 1 | 72 | 26962 - 27033 (+) | AAGTAATTAACTTAGAGGTAAAGTTTCTGCTTTACATACAGAAG ACCATTGGTTCGAATCCAATATTACTTA |
| **22.** | PF3D7_API05300 | Arginine | TCT | 1 | 72 | 27035 - 27106 (+) | AAATCTATAATTTAATGGATAAAATAAAAACCTTCTAAGTTTTA TATGTAAGTTCAAATCTTACTAGATTTA |
| **23.** | PF3D7_API05500 | Asparagine | GTT | 1 | 72 | 27203 - 27274 (-) | TTCTTAATAGCTTAGTGGTTAAAGCATTCGGCTGTTAACCGAAA TACACTAGTTCAATTCTAGTTTAAGAAG |
| **24.** | PF3D7_API05600 | Alanine | TGC | 1 | 71 | 27304 - 27374 (+) | GGGAATATAGTTTAATGGTAAAATCTTATTTTTGCATAATAAAG ATAGTAGTTCAATTCTACTTATTTCCA |
| **25.** | PF3D7_API05800 | Isoleucine | GAT | 1 | 72 | 28994 - 29065 (+) | ATAGGTTTTTAGTTTAATGGTTAAAACATACTCTTGATAAGGGT AAAATTTTAGTTCAATTCTAAAATAACC |
| **26.** | PF3D7_API03100 | Phenylalanine | GAA | 1 | 72 | 9873 - 9944 (-) | GTCATAATAGCTCAATGGTAGAGCAATGGATTGAAGATCCATGT GTTATCAGTTCAAATCTGATTTTTGACA |
| **27.** | PF3D7_API06000 | Alanine | TGC | 1 | 71 | 30816 - 30886 (-) | GGAATATAGTTTAATGGTAAAATCTTATTTTTGCATAATAAAGA TAGTAGTTCAATTCTACTTATTTCCAT |
| **28.** | PF3D7_API06100 | Asparagine | GTT | 1 | 72 | 30917 - 30988 (+) | TTCTTAATAGCTTAGTGGTTAAAGCATTCGGCTGTTAACCGAAA TACACTAGTTCAATTCTAGTTTAAGAAG |
| **29.** | PF3D7_API06300 | Arginine | TCT | 1 | 72 | 31085 - 31156 (-) | AAATCTATAATTTAATGGATAAAATAAAAACCTTCTAAGTTTTA TATGTAAGTTCAAATCTTACTAGATTTA |
| **30.** | PF3D7_API06400 | Valine | TAC | 1 | 68 | 31162 - 31229 (-) | AAGTAATTAACTTAGAGGTAAAGTTTCTGCTTTACATACAGAAG ACCATTGGTTCGAATCCAATATTA |

| | | | | | | | |
|---|---|---|---|---|---|---|---|
| **31.** | PF3D7_API06500 | Arginine | ACG | 1 | 71 | 31251 - 31321 (+) | AACTTGTAATTTAATGGATAAAATATATAAATACGAATTATAAA ATAAAAGTTCAATTCTTTTCAAGTTTA |
| **32.** | PF3D7_API06600 | Methionine | CAT | 1 | 75 | 31327 - 31401 (+) | AGCGAAATAGAGCATAAGGAAAGTTCGTCGGATTCATGCTCCGA AGGTAATCGGTTCAATTCCGTTTTTCGCTTA |
| **33.** | PF3D7_API06800 | Threonine | TGT | 1 | 73 | 34153 - 34225 (+) | GCTAAAGTAGCTTAATTGGTAAAGCAACTGATTTGTAATCAGTA GATTATGAGTTCAAATCTCACCATTAGCT |

**Table S2:** List of mature tRNAs (non-coding RNA) present in the nucleus of *Plasmodium falciparum* 3D7 retrieved from PlasmoDB Database and verified through tRNAscan-SE 2.0. Nucleotide sequence not found in tRNAscan-SE 2.0 are highlighted in yellow and their transcript length is represented in blue.

| S.No. | PlasmoDB ID | tRNA type | Anticodon loop (5′-3′ direction) | Transcript length | Genomic Location/ Chromosome Number | Sequence (Genomic) |
|---|---|---|---|---|---|---|
| **1.** | PF3D7_1103300 | Leucine | CAA | 99 (84) | 144841 - 144939(-) Chromosome 11 | GCACGGATGGCTGAGTGGTCTAAAG CGTCAGACTCAAGATCTGATGAACG TAAGTTCGCATGGGTTCGAACCCCA TTTCGTGCATATTTAAAAAGGCCA |
| **2.** | PF3D7_0620900 | Leucine | CAG | 81 | 861685 - 861765(-) Chromosome 6 | GACAGAATGGCCGAGCGGTCTAAGG CGCTACAGTCAGATTGTAGTCTGTA TAAGGCGTGGGTTCGAATCCCGCTT CTGTCA |
| **3.** | PF3D7_0510600 | Leucine | TAG | 80 | 451355 - 451434(+) Chromosome 5 | GTCAGGATGGCCGAGTGGTCTAAGG CGCAGCGTTTAGGCCGCTGTCCGTA AGGGCGTGGGTTCGAACCCCACTCC TGACA |
| **4.** | PF3D7_0527800 | Leucine | TAA | 83 | 1151227 - 1151309(-) Chromosome 5 | GCACGGGTGCCCGAGTGGTTAAGGG GGTGGACTTAAGATCCTCTGGTCAC TAGACCGCGTGGGTTCGAACCCCAC CTCGTGCA |
| **5.** | PF3D7_0714800 | Leucine | AAG | 80 | 671656 - 671735(+) Chromosome 7 | GACAGAATGGCCGAGTGGTCTAAGG CGCAACGTTAAGGCCGTTGTCCGAA |

| | | | | | | |
|---|---|---|---|---|---|---|
| | | | | | | AGGGCGTGGGTTCGAACCCCACTTC TGTCA |
| **6.** | PF3D7_1252000 | Glutamine | TTG | 72 | 2113730 - 2113801(-) Chromosome 12 | GGTTTCGTAGTGTAGTGGTTAGCAC TGAGGACTTTGAATCCTCCAACCCG GGTTCGAGTCCCGGCGAGACCT |
| **7.** | PF3D7_0203500 | Glutamine | CTG | 72 | 164269 - 164340(+) Chromosome 2 | GGTTCTGTAGTGTAGTGGTTAGCAC TGCAGACTCTGACTCTGCAAACCTG GGTTCAAATCCCAGCAGAACCT |
| **8.** | PF3D7_1103200 | Glycine | TCC | 73 (72) | 143353 - 143425(+) Chromosome 11 | GGCGTCAATAGTCTAACGGCCATGA TACCTGCCTTCCAAGCAGGTGACCC GGGTTCGACTCCCGGTTGACGCA |
| **9.** | PF3D7_1370200 | Glycine | GCC | 71 | 2783766 - 2783836(+) Chromosome 13 | GCATCTGTGGTCTAGTGGTAGAATA CTTCGTTGCCATCGAAGTGACCCGG GTTCGATTCCCGGCAGATGCA |
| **10.** | PF3D7_0411600 | Glutamic acid | CTC | 72 | 518154 - 518225(-) Chromosome 4 | TCCCACGTGGTCTAGTGGCTAGGAT ATTCGGCTCTCACCCGAAAGGCCCG GGTTCAATTCCCGGCGTGGGAA |
| **11.** | PF3D7_0527700 | Glutamic acid | TTC | 73 | 1150234 - 1150306(+) Chromosome 5 | TCCCATGTAGTCTAGGCGGTTAGGA TATTCGGCTTTCACCCGAACGACCC GGGTTCGAGTCCCGGCGTGGGAA |
| **12.** | PF3D7_0529600 | Arginine | TCG | 73 | 1208390 - 1208462(+) Chromosome 5 | GACGGCGTAGCCTAATGGATAAGGC GTCGGTCTTCGGAACCGAAGATTGC GGGTTCGAGTCCCGTCGTCGTTA |
| **13.** | PF3D7_1370000 | Arginine | CCT | 74 | 2781622 - 2781695(+) Chromosome 13 | GCACCAGTGGCGTAATTGGATAGCG CAATGCCTTCCTAAGGCAAAGGTTA TGGGTTCGAGTCCCATCTGGTGTA |
| **14.** | PF3D7_1369800 | Arginine | ACG | 74 | 2780354 - 2780427(+) Chromosome 13 | GGGCCGGTAGTTCAGTTGGATAGAA TGCCCGACTACGGATCGGGAGGTCG TGGGTTCGACTCCTGCCCGGCCTA |
| **15.** | PF3D7_1341000 | Arginine | TCT | 74 | 1631333 - 1631406(+) Chromosome 13 | GCCTCTGTGGCGCAATTGGATAGCG CGTTGGACTTCTAATCCAAAGGCTG CGGGTTCGAATCCCGCCAGGGGTA |
| **16.** | PF3D7_1418400 | Proline | AGG | 72 | 778092 - 778163(+) Chromosome 14 | GGCTACTTAGTCTAGTGGTATGATT CTCTCTTAGGGTGGGAGAGGTCCCG GGTTCGATTCCCGGAGTAGCCC |

| | | | | | | |
|---|---|---|---|---|---|---|
| **17.** | PF3D7_1216800 | Proline | CGG | 72 | 666884 - 666955(+) Chromosome 12 | GGCTACTTGATCTAGTGGTATGATT CTTGCTTCGGGTGCAAGACGTCCCG GGTTCGATTCCCGGAGTAGCCC |
| **18.** | PF3D7_1339200 | Proline | TGG | 72 | 1576926 - 1576997(-) Chromosome 13 | GGCTACTTAGTCTAGTGGTATGATT CTCGCTTTGGGTGCGAGAGGTCCCG GGTTCAATTCCCGGAGTAGCCC |
| **19.** | PF3D7_1438300 | Methionine | CAT | 74 | 1546224 - 1546297(+) Chromosome 14 | GGGTAATTGGCGCAGTTGGTTAGCG CGCGGGTCTCATAATCCCGAGGTCG TGAGTTCGATCCTCACATTACCCA |
| **20.** | PF3D7_1339100 | Methionine | CAT | 72 | 1575356 - 1575427(+) Chromosome 13 | AGCAGCGTAGCTCAGAGGAAGAGTG GGGGGCTCATAACCCCCAGGACCGT GGATCGAAACCACGCGCTGCTA |
| **21.** | PF3D7_0730700 | Threonine | TGT | 72 | 1319011 - 1319082(-) Chromosome 7 | GCCACCTTAGCACAGTGGTAGTGCG TTGGTCTTGTAAACCAAAGGTCGGG AGTTCGATCCTCCCAGGTGGCT |
| **22.** | PF3D7_0706800 | Threonine | AGT | 73 | 326072 - 326144(+) Chromosome 7 | GCCTGCATAGCTTAGTTGGTAGAGC ATCCGCCTAGTAAGCGGAAGGTCAT CGGTTCGACTCCGGTTGTAGGCT |
| **23.** | PF3D7_1355400 | Threonine | CGT | 73 | 2197906 - 2197978(-) Chromosome 13 | GCCATCTTAGCACAGTTGGTAATGC GTTTGATTCGTAATCAAAAGATCGT GGGTTCGACTCCCACAGGTGGCT |
| **24.** | PF3D7_1251900 | Valine | AAC | 73 | 2112878 - 2112950(+) Chromosome 12 | GCGGGCATGGTCTAGTGGCTATGAC GCCTGCCTAACACGCAGGAGATCCC GAGTTCGATCCTCGGTGCCCGTA |
| **25.** | PF3D7_0312600 | Valine | CAC | 77 (73) | 531213 - 531289 (+) Chromosome 3 | GCGAGCATGGTCTAGTGGCTATGAC GTTCGCCTCACACGCGAAAGATCCC GAGTTCGATCCTCGGTGCTCGTATT AA |
| **26.** | PF3D7_0730600 | Valine | TAC | 73 | 1318059 - 1318131(+) Chromosome 7 | GCGGGCATGGTCTAGTGGCTATGAC GCCTGCCTTACACGCAGGAGATCCC GAGTTCGATCCTCGGTGCCCGTA |
| **27.** | PF3D7_1337600 | Serine | CGA | 82 | 1524111 - 1524192(-) Chromosome 13 | GACAGTTTGTCCGAGTGGTTAAGGA GGTTGACTCGAAATCAACTGGGCTC TGCCCGCACAGGTTCAAATCCTGTA GCTGTCG |

| | | | | | | |
|---|---|---|---|---|---|---|
| **28.** | PF3D7_0410100 | Serine | AGA | 82 | 469218 - 469299(+) Chromosome 4 | GACAGTGTGTCCGAGTGGTTAAGGAGTCAGACTAGAAATCTGGTAGGCTTTGCCTGCGCAGGTTCGAATCCTGCCGCTGTCG |
| **29.** | PF3D7_0621600 | Serine | TGA | 82 | 883563 - 883644(-) Chromosome 6 | GACAGTTTGCCCGAGTGGTTAAGGGGTTGGACTTGAAATCCAATGAGCTTTGCTTGCGCAGGTTCGAATCCTGCAGCTGTCG |
| **30.** | PF3D7_0714900 | Serine | GCT | 82 | 672735 - 672816(-) Chromosome 7 | GATAACGTGCCCGAGTGGTTAAGGGGTTGGACTGCTAATCCAATGGGTTCTGCCCGCGCAGGTTCGATTCCTGCCGTTGTCG |
| **31.** | PF3D7_0707000 | Lysine | CTT | 73 | 327865 - 327937(+) Chromosome 7 | GCTGCCTTAGCTCAGTCGGTAGAGCGCCAGACTCTTAATCTGGTGGTCAGGGGTTCGAGCCCCCTAGGCAGCT |
| **32.** | PF3D7_0707100 | Lysine | TTT | 73 | 329012 - 329084(-) Chromosome 7 | GCCTCTTTAGCTCAGTTGGTAGAGCGCCAGACTTTTAATCTGGTGGTCAGGGGTTCGAGTCCCCTAGGAGGCT |
| **33.** | PF3D7_0312700 | Isoleucine | AAT | 76 (74) | 531898 - 531973(-) Chromosome 3 | GGTCCTATAGCTCAGTTGGTTAGAGCGTACGGCTAATAACCGTAAGGTCGGCGGTTCGAGACCGCCTGGGACCAAA |
| **34.** | PF3D7_0410200 | Isoleucine | TAT | 73 | 469858 - 469930(-) Chromosome 4 | GATCCATTAGCTCAGTGGTTAGAGCGTCGGTCTTATGTACCGAAGGTCGTGGGTTCGAAACCCACATGGATCA |
| **35.** | PF3D7_0620800 | Alanine | CGC | 72 | 861017 - 861088(+) Chromosome 6 | GGGCTAGTAGTGTAGTGGTATCACGTTCGCTTCGCATGCGAAAGTTCCCGGGTTCGATTCCCGGCTGGTCCA |
| **36.** | PF3D7_0702700 | Alanine | AGC | 73 | 111473 - 111545(-) Chromosome 7 | GGGCGACTAGCTCAAGTGGTAGAGCGCTCGCTTAGCATGCGAGAGGTACGGGGATCGATACCCCGGTCGTCCA |
| **37.** | PF3D7_0411500 | Alanine | TGC | 72 | 517722 - 517793(+) Chromosome 4 | GGGCAGGTGGTGTAGTGGTATCACGCTTGATTTGCATTCAAGAGGTCCGGGGTTCAATTCCCCGTCTGTCCA |

| | | | | | | |
|---|---|---|---|---|---|---|
| **38.** | PF3D7_0403000 | Asparagine | GTT | 74 | 174829 - 174902(+) Chromosome 4 | GGTTCCGTAGCTCAGTTGGTTAGAG CGTGCGGCTGTTAACCGCAAGGTCG TTGGTTCGATCCCAGCCGGTACCG |
| **39.** | PF3D7_0714700 | Asparagine (ASP) | GTC | 72 | 670795 - 670866(+) Chromosome 7 | TCCGAGATAGTATAGTGGCAAGTAT TTCCGCCTGTCACGCGGAAGACCCG GGTTCAATTCCCGGTCTCGGAG |
| **40.** | PF3D7_1370100 | Cysteine | GCA | 72 | 2783150 - 2783221(-) Chromosome 13 | GGGCGTGTAGCTCAGCGGTAGAGCA GCTGACTGCAGATCAGCGGGTCCAT GGTTCAAATCCGTGCGCGCCCT |
| **41.** | PF3D7_0706900 | Histidine | GTG | 72 | 326726 - 326797(-) Chromosome 7 | GTCCAAATCGTCTAGTGGTTAGGAC TCCACGCTGTGGACGTGGCAACGTA GGTTCGAATCCTGCTTTGGACA |
| **42.** | PF3D7_0514400 | Phenylalanine | GAA | 73 | 603951 - 604023(-) Chromosome 5 | GCCGTGATAGCTCAGTTGGGAGAGC GTCAGACTGAAGATCTGAAGGTCCC TGGTTCGATCCCTGGTCACGGCA |
| **43.** | PF3D7_1369900 | Tryptophan | CCA | 72 | 2781080 - 2781151(-) Chromosome 13 | GGGCCTATGGCGCAACGGTAGCGCG TCTGACTCCAGATCAGAAGGCTGGG GGTTCGAATCCCTCTGGGCTCA |
| **44.** | PF3D7_0702800 | Tyrosine | GTA | 84 (73) | 112049 - 112132(+) Chromosome 7 | CCGATGATAGCTCAGTTGGTAGAGC GGCAGACTGTAGttgaaatggttAT CTGTTGGTCACCGGTTCGATTCCGG TTCATCGGA |
| **45.** | PF3D7_1438200 | Selenocysteine | TCA | 90 | 1545146 - 1545235(-) Chromosome 14 | GCACCGATGAGTTAGCATGGTTGCT AAGTATGACTTCAAATCATTTGGCG TAGTTTTTCTGCGCAGAGGTTCGAT TCCTCCTTCGGTGCG |

**Table S3: Primer sequences for the real-time PCR analysis of *Pf*tRNA$^{Leu}$ (Apicoplast).**

| Genes | Primer sequences (5′-3′) |
|---|---|
| *Pf*tRNA$^{Leu}$ (Nuclear) | FP: 5′-ATA ggA TCC TAA TAC gAC TCA CTA TAg ggA ATT TCA TAT gTT TTA TTA Ag-3′<br>RP: 5′-CgA CTC gAA TTC TAA ATT TTT TTT ATT CAA TgT AC-3′ |
| *Pf*tRNA$^{Leu}$ (Apicoplast) | FP: 5′-ATA ggA TCC TAA TAC gAC TCA CTA TAg ggA gAg ATA Tgg TgA AAT TTg-3′<br>RP: 5′-gAC TCg AAT TCT AgA gAT AAA ggg ATT TgA AC-3′ |
| 18s rRNA | FP: 5′-CCGCCCGTCGCTCCTACCG-3′<br>RP: 5′-CCTTGTTACGACTTCTCCTTCC-3′ |

```
P.falciparum 3D7            -AGAGATATGGTGAAATTTGGTATACACAATGGACTTAAAATAATTTGAG
P.falciparum strain GB4     -AGAGATATGGTGAAATTTGGTATACACAATGGACTTAAAATAATTTGAG
P.falciparum strain HB3     -AGAGATATGGTGAAATTTGGTATACACAATGGACTTAAAATAATTTGAG
Plasmodium vivax            -AGAGATATGGTGAAATTAGGTATACACAATGGACTTTATTAAATATTAA
Plasmodium yeolii           TAGAGATAAAG-GGACTCGAACCCTTACAAC-------AATTATTGTTAA
Plasmodium strain IT        TAGAGATAAAG-GGATTTGAACCCTTACAAC-------AAT-AATGTTAA
                             *******  * * * *         ****        *   * * * *

P.falciparum 3D7            TTAAT----TATTA-TTAATATTAAATTTTTAAGAAAATAT----ATAAT
P.falciparum strain GB4     TTAAT----TATTA-TTAATATTAAATTTTTAAGAAAATAT----ATAAT
P.falciparum strain HB3     TTAAT----TATTA-TTAATATTAAATTTTTAAGAAAATAT----ATAAT
Plasmodium vivax            TAAATATTATATGAGTTAATTATAAATTAATAATAGTATTTTATAAGAAA
Plasmodium yeolii           TGGATTTT--TCTAATTAAATTTAGACTTTTTATAAAAATATATACATAAA
P.falciparum strain IT      TGGATTT--TCTTAATAAATTTAGACTTTTTATAAA----------TAAA
                            *  **    * * * * ***    * **  **  *            **

P.falciparum 3D7            ATATTTT-----TTTAAATTCTGTAATATATTTTAAAA-----------T
P.falciparum strain GB4     ATATTTT-----TTTAAATTCTGTAATATATTTTAAAA-----------T
```

```
P.falciparum strain HB3      ATATTTT-----TTTAAATTCTGTAATATATTTTAAAA-----------T
Plasmodium vivax             ATAATTT-----TTTAAATTCTGTAATATATTAT---------------T
Plasmodium yeolii            GTCGTTTGAATATATAACTAATATATTACAGAATAAAAATTATTTTTTCT
P.falciparum strain IT       GTCTTTTGAATATATAAATTTTAAAATATATTACAGAA-----------T
                              *  ***     * *** *  *  * ** *                   *

P.falciparum 3D7             TTATATATTCAAAAGACTTT---ATTTATAAAAAGTCTAAATTTATTAA-
P.falciparum strain GB4      TTATATATTCAAAAGACTTT---ATTTATAAAAAGTCTAAATTTATTAA-
P.falciparum strain HB3      TTATATATTCAAAAGACTTT---ATTTATAAAAAGTCTAAATTTATTAA-
Plasmodium vivax             TTATATATTCAAAAGACTTTTTGATTTTTTAAAAGTCTAAATTTAAAAA-
Plasmodium yeolii            TTATATATTAAAATTATTAAT--TTATTTATAAAATTAACTCATAACATA
P.falciparum strain IT       TTAAAAAAATATATTATATAT--TTTCTTA-AAAATTTAATATTAATA--
                             *** * *   * *  *        *   *  *** *  *    **  *

P.falciparum 3D7             --GAAAATCCATTAACATTA-TT-------GTTGTAAGGGTTCAAATCC-
P.falciparum strain GB4      --GAAAATCCATTAACATTA-TT-------GTTGTAAGGGTTCAAATCC-
P.falciparum strain HB3      --GAAAATCCATTAACATTA-TT-------GTTGTAAGGGTTCAAATCC-
Plasmodium vivax             --GAAAATCCATTAACAATAATT-------GTTGTAAGGGTTCAAATCC-
Plasmodium yeolii            ATATAACTATTATAATATTATTTAAAGTCCATTGTGTATACCAAATTTCA
P.falciparum strain IT       --ATAATTAACTCAA-ATTATTTTAAGTCCATTGTGTATACCAAATTTCA
                                 ** *     ** * ** **        ****        ** * *



P.falciparum 3D7             CTTTATCTCTA
P.falciparum strain GB4      CTTTATCTCTA
P.falciparum strain HB3      CTTTATCTCTA
Plasmodium vivax             CTTTATCTCTA
Plasmodium yeolii            CCATATCTCT-
P.falciparum strain IT       CCATATCTCT-
                             *  *******
```

**Fig. S1:** Multiple sequence alignment of pre-tRNA$^{leu}$ (apicoplast) of *Plasmodium falciparum 3D7* with its homologs in different *Plasmodium* strains (GB4, HB3, IT) and species (*Plasmodium vivax*, *Plasmodium Yeolii*, showing its conserved nature.

```
Thermoplasma acidophilum                 HSASQVDMVYSDLVGRGCIVKTGFKYGANFRVYLGRD---------SQHAEYLVSVMPEE   247
PfTSEN1 (PF3D7_1248000)                  NKFVVK--YKEYFEENSFIVKKGDIYGADFLLYITEQ--------KYAHSLYAVYIIYKH   174
Hepatocystis sp. ex P. tephrosceles      LKLLKK--CKKYFKNNGYIIKKGDLYGGDYLLYLTNE--------KYTHSVYVVYFIKKE   185
Saccharomyces cerevisiae (SEN34)         AGKMQTYFLYKALRDQGYVLSPGGRFGGKFIAYPGDP--------LRFHSHLTIQDAI-D   227
Arabidopsis Thaliana (ATSEN1)            QDFAILYKAYSHLRSKNWIVRSGLQYGVDFVVYRHHP--------SLVHSEYAVLVQSIG   167
Arabidopsis Thaliana (ATSEN2)            PNFPMFFKAYSHLRSKNWVLRSGLQYGVDFVAYRHHP--------SLVHSEYSVLVQS-G   166
Candidatus M. acidiphilum                TDIMKLYDVYKDWRTKGYVVKTGFKFGTNFRIYFPGAKPIKENN-EWIHSKHVLHVFPRD   262
Saccharomyces Cerevisiae (SEN2)          HSFVRSYVIYHHYRSHGWCVRSGIKFGCDYLLYKRGP--------PFQHAEFCVMGLDHD   308
Mus musculus (SEN2)                      PTFRTTYMAYHYFRSKGWVPKVGLKYGTDLLLYRKGP--------PFYHASYSVIIELLD   288
Homo sapiens (SEN2)                      PTFRTTYMAYHYFRSKGWVPKVGLKYGTDLLLYRKGP--------PFYHASYSVIIELVD   388
Mus musculus (SEN34)                     PAHELRYSIYRDLWERGFFLSAAGKFGGDFLVYPGDP--------LRFHAHYIAQCWSAE   272
Homo sapiens (SEN34)                     PAHELRYSIYRDLWERGFFLSAAGKFGGDFLVYPGDP--------LRFHAHYIAQCWAPE   266
Archaeoglobus fulgidus                   RNFDRRYEVYRNLKERGFVVKTGFKFGSEFRVYRKVE-----SVDDLPHSEYLVDIA-DS   267
PfTSEN2 (PF3D7_1454100)                  RKREI---VFHDL-NKLFIVVDGFKYGADFVLYKNNV--------DEEHGFALVFIKEEN   53
Nanoarchaeum equitans                    ERFLIRYKAYKELRDKGYTLGTALKFGADFRVYDIGVIPKKGKRSEREHSKWVLYPVSKD   113
Sulfolobus tokodaii                      PRFRILYSVYEDLREKGYVVRSGIKYGADFAVYTIGP--------GIEHAPYLVIALDEN   136
Sulfolobus solfataricus                  NKFEILYKVYEDLREKGFIVRSGVKYGADFAVYTLGP--------GLEHAPYVVIAVDID   138
Aeropyrum pernix                         PRFSMLYNIYRDLRERGFVVRSGLKFGSDFAVYRLGP--------GIDHAPFIVHAYSPE   144
Pyrobaculum aerophilum                   RNFDEIYKIYKYFRDLGYVVKSGLKFGALFSVYEKGP--------GIDHAPMVVVFLEPD   139
                                                               .  :*     *              *.

Thermoplasma acidophilum                 E----------RWYSISRGVRVASSVRKTMIYASIYKNE--VRY----------VALKRV   285
PfTSEN1 (PF3D7_1248000)                  Y----------ILRDLIRILRVSHSIKKVILILWNELNCFNIS----DTLVYIKSYKYK   220
Hepatocystis sp. ex P. tephrosceles      K----------TLRELIQILRLSCSIKKVIIVLETDTIPLDIT----HSMVFIETYFYK   231
Saccharomyces cerevisiae (SEN34)         Y-----HNEPIDLISMISGARLGTTVKKLWVIGGVAEETKETHF----------FSIEWA   272
Arabidopsis Thaliana (ATSEN1)            G-----NDRLKVWSDIHCSVRLTGSVAKSLLVLYVNRKVNT-------EKMNLPLCLEDY   215
Arabidopsis Thaliana (ATSEN2)            D-----SDRLRVWSDIHCAVRLSGSVAKTLLTLYVNGNFKG-------EDVNLLVCLENF   214
Candidatus M. acidiphilum                S--------KLIISEWARAIRVAHSVRKTFILAIPGKTRKKKLA----------IDFELY   304
Saccharomyces Cerevisiae (SEN2)          V--------SKDYTWYSSIARVVGGAKKTFVLCYVERLISEQEAIALWKSNNFTKLFNSF   360
Mus musculus (SEN2)                      DNYEGSLRRPFSWKSLAALSRVSGNVSKELMLCYLIKPSTMTAE-----DMETPECMKRI   343
Homo sapiens (SEN2)                      DHFEGSLRRPLSWKSLAALSRVSVNVSKELMLCYLIKPSTMTDK-----EMESPECMKRI   443
Mus musculus (SEN34)                     D--------PIPLQDLVSAGRLGTSVRKTLLLCSP-QPDGKVVY----------TSLQWA   313
Homo sapiens (SEN34)                     D--------TIPLQDLVAAGRLGTSVRKTLLLCSP-QPDGKVVY----------TSLQWA   307
Archaeoglobus fulgidus                   R--------EIRLIDLARAVRLAQNVRKRMVFAYGK------NY----------LCFERV   303
PfTSEN2 (PF3D7_1454100)                  I--------CLNEKEKNIIVRICESVKKKAIIAYINEENKSIRY----------EEVFRK   95
Nanoarchaeum equitans                    E--------TFDFYEFASKNRVAHSTRKKMLMGIVS---DKIEF----------IEVSWK   152
Sulfolobus tokodaii                      S--------QISSNEILGFGRVSHSTRKELILGIVNLTNGKIRY----------IMFKWL   178
Sulfolobus solfataricus                  E--------EITPHELLSFGRVSHSTKKRLVLALVDRKSEGIRY----------IMFKWV   180
Aeropyrum pernix                         D--------NIDPVEIVRAGRLSHSVRKKFVFAV--TRGGDVSY----------LMIDWF   184
Pyrobaculum aerophilum                   K--------GISATDITRGGRLSHSVRKTWTLATVLRQTGEVVL----------LGFGWA   181
                                                             *:     *
```

**Fig. S2:** Multiple sequence alignment of *Pf*TSEN1 with its homologs in other species. Residues His, Lys and Tyr involved in the formation of the catalytic triad and playing role in activity of *Pf*TSEN1 are marked in yellow color.

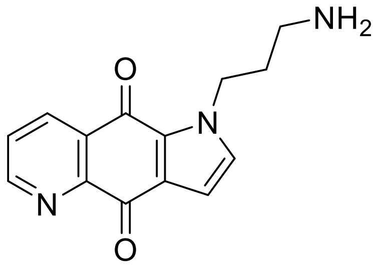


N1- (2,2-dimethoxyethyl) propane-1,3-diamine (1.57 g, 9.7 mmol, 1.5 eq.) was added to a solution of quinoline-5,8-dione 19 (1.03 g, 6.5 mmol, 1 eq.) and Cerium (III) chloride heptahydrate (0.16 g, 0.4 mmol) in 40 mL dry acetonitrile, the resulting reaction mixture was treated for 24 hours at room temperature in an ultrasonic bath. The resulting reaction mixture was used for the further synthesis without further work-up.

Aqueous sulfuric acid (5 M, 7 mL) was added to the reaction mixture and heated to 70 °C for 24 hours while stirring. Subsequently, acetonitrile was removed under reduced pressure, DCM was added to the aqueous solution and the resulting reaction mixture filtered. The filtrate was then washed with DCM, made basic with 4 M sodium hydroxide solution and extracted three times with DCM. The solvent from the combined organic phases was removed under reduced pressure. The desired product was found as a light brown solid (0.55 g, 2.2 mmol, 33.8%).

1H-NMR (500 MHz, CDCl3) δ [ppm] = 8.85 (dd, 1 H, J = 3.3, 1.3 Hz, -CHAr -), 8.33 (dd, 1 H, J = 6.7, 1.3 Hz, -CHAr -), 7.56 (t, 1 H, J = 3.1 Hz, -CHAr-), 7.40 (d, 1 H, J = 5.4 Hz, -CHAr-), 7.09 (d, 1 H, J = 4.9 Hz, -CHAr -), 4.41 (m, 2H, NH2-CH2-CH2-CH2), 2.64 (t, 2 H, J = 2.9 Hz, NH2 -CH2 -CH2 -CH2), 2.14 (m, 2 H, NH2 -CH2 -CH2 -CH2)

13C-NMR (125 MHz, CDCl3) δ [ppm] = 172.51 (Cq=O), 162.44 (Cq=O), 153.49 (CH), 148.29 (Cq), 138.63 (Cq), 136.22 (CH), 132.38 (Cq), 128.95 (CH), 127.66 (Cq), 122.04 (CH), 102.51 (CH), 46.43

(CH2 ), 40.27 (CH2 ), 29.65 (CH2)

**Fig. S3: Schematic representation for TSEN$_i$ synthesis.**

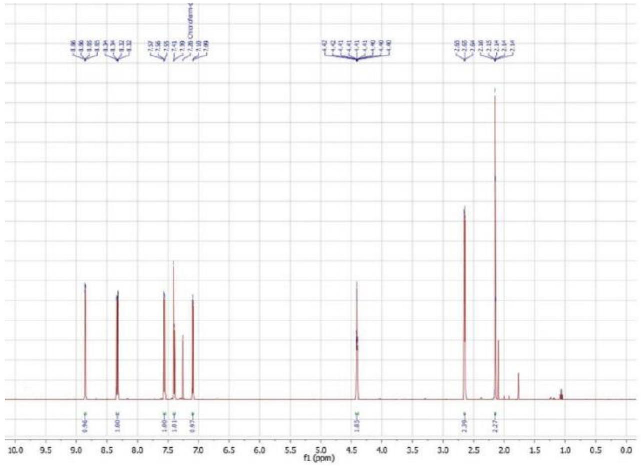


**Fig. S4: $^{1}$H NMR spectrum of TSEN$_{i}$.**

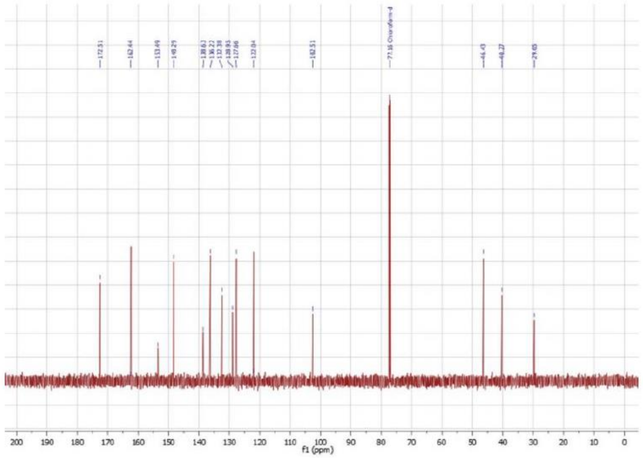


**Fig. S5: $^{13}$C-NMR spectrum of TSEN$_{i}$.**
